# Viable and efficient electroporation-based genetic manipulation of unstimulated human T cells

**DOI:** 10.1101/466243

**Authors:** Pinar Aksoy, Bülent Arman Aksoy, Eric Czech, Jeff Hammerbacher

## Abstract

Electroporation is the most feasible non-viral material delivery system for manipulating human T cells given its time- and cost-effectiveness. However, efficient delivery requires electroporation settings to be optimized for different devices, cellular states, and materials to be delivered. Here, we used electroporation to either induce exogenous gene expression in human primary T cells by plasmids or in vitro transcribed (IVT) mRNA and also target endogenous genes by Cas9 ribonucleoproteins (RNPs). We characterized the electroporation conditions both for activated and unstimulated human T cells. Although naive cells are non-dividing and therefore their genetic manipulation is harder compared to activated T cells, we developed the technical ability to manipulate both naive and memory cells within the unstimulated T cell population by IVT mRNA and Cas9 RNP electroporation. Here, we outline the best practices for achieving highly-efficient genetic manipulation in primary T cells without causing significant cytotoxicity to the cells. Because there is increasing evidence for “less-differentiated” T cells to have better anti-tumor activity for immunotherapy, manipulating naive T cells with high efficiency is also of high importance to clinical applications and to study the biology of these cells.

## Introduction

Adoptive cell transfer (ACT) is an immunotherapy method in which a cancer patient’s own or allogeneic immune cells (*e.g.* T cells) are infused back to the patient. These cells can be genetically edited to improve their anti-tumor activity. Because T cells in general, and naive T cells specifically, have been challenging to genetically manipulate, activation of T cells has been a prerequisite for T cell engineering for clinical applications. However, activation of the cells pushes them towards their differentiation program and the longer the cells are cultured *ex vivo* to achieve a certain number, the more exhausted they become. Several studies have shown superior anti-tumor effect of “less-differentiated” cells in ACT–*i.e.* naive cells do better than memory cells and central memory (CM) cells do better than effector memory cells (EMs) (Gattinoni et al., 2005; Hinrichs et al., 2011, 2009). Therefore, developing the techniques for genetic manipulation of naive cells with high efficiency and viability is important for these applications. Aside from the clinical applications, achieving naive T cell manipulation is also important for studying the biology of these cells with minimal perturbation to their unstimulated state.

For most cell types, genetic manipulation is achieved by transfection. However, transfection of T cells through commonly-used transfection reagents has not been possible due to high toxicities associated with the reagents, such as lipofectamine (Ebert et al., 1997). Another way of delivering materials into cells is by electroporation– *i.e.* opening pores on the cell membrane. Electroporation has been widely used since its first introduction in 1982 (Neumann et al., 1982). In recent years, relatively more efficient electroporation devices have been made commercially available (e.g. Lonza’s nucleofector or Thermo Fisher’s Neon electroporation devices). The first published study to show plasmid electroporation in unstimulated human T cells achieved 37% efficiency and 32% viability (Bell et al., 2001). In the same study, Bell et al also showed that in 24 hours, the frequency of both GFP-expressing cells and viable cells declined compared to 7 hours post-electroporation. An earlier study using phytohemagglutinin (PHA)-activated human T lymphocytes resulted in very low transgene expression (15%) (Van Tendeloo et al., 2000). In 2011, a broad optimization study using the Neon electroporation machine showed 59.6% efficiency and 34.6% viability in unstimulated CD8+ T cells (Liu et al., 2011). Later, in 2013, another group showed that CD3/CD28-activated T cells were vulnerable to plasmid electroporation by nucleofection and because of this, plasmid electroporation in activated cells was not achieved (Chicaybam et al., 2013). The same study showed ∼45% electro-transfection efficiency and 25% viability in unstimulated PBMCs. They also showed that when PBMCs were activated 24 hours after plasmid electroporation, GFP expression frequencies remained higher than 30% for 7 days (Chicaybam et al., 2013). A more recent paper from 2018 showed that plasmid electroporation could yield 40% efficiency in CD3/CD28 Dynabead-activated human T cells, however, it also concluded that unstimulated cells could not be efficiently electroporated with plasmids (<5% efficiency) (Zhang et al., 2018).

Studies from the 2000s investigated mRNA electroporation of PBMCs with contradicting results. One paper claimed that both unstimulated and CD3-stimulated T cells could be efficiently electroporated with GFP mRNA (Zhao et al., 2006). An earlier paper concluded that PHA-stimulated T cells could efficiently be electroporated with GFP mRNA, however unstimulated PBMCs could not (Smits et al., 2004). The most recent paper on RNA electroporation of unstimulated CD8+ T cells described a double sequential electroporation method to knock down endogenous TCRs and then insert a tumor-specific TCR mRNA (Campillo-Davo et al., 2018). Another set of papers showed successful gene knockouts by Cas9 RNPs in both unstimulated and activated cells. In 2015, Marson Lab reported successful utilization of Cas9 ribonucleoproteins (RNPs) for gene editing in activated human T cells (Schumann et al., 2015). However, editing unstimulated cells has remained a challenge for the last couple of years. A paper from 2018 was the first to show efficient knockout in both human and mice unstimulated T cells using Cas9 RNPs (Seki and Rutz, 2018). In this 2018 paper, the group optimized the buffers and electroporation settings using Lonza’s nucleofector and, most importantly, showed that a combination of 3 sgRNAs increased target gene knockout efficiency compared to a single-gRNA-mediated-targeting.

Here, we electroporated both activated and unstimulated T cells, which were isolated from healthy human donors, with plasmids, mRNA, or Cas9 RNPs. Although successful electroporation of unstimulated cells by these materials have been shown before by others as previously mentioned, to our knowledge, we are the first to show the efficiencies within the subpopulations of unstimulated cells and therefore clearly show our ability to manipulate not only memory cells (CM and EM) but also naive T cells. We also applied our knowledge to mouse T cells and electroporated them with murine TCRs. After showing that these TCR-electroporated mouse T cells were cytotoxic against cognate antigen-presenting cells, we also showed that these murine TCRs, when electroporated into human T cells, could make human T cells cytotoxic as well.

## Results

### Electroporation of plasmids into activated and unstimulated T cells

#### Plasmid electroporation into activated cells

To assess electroporation as a technique for genetic manipulation of human T cells, we activated the cells with CD3/CD28 antibody-coated beads. On the second day of activation, we debeaded the cells and electroporated them with a GFP plasmid containing the PEST domain and a nuclear localization signal (NLS). The cells were electroporated at a concentration of 7.5 ug DNA per million cells. The next day, the frequency of GFP+ cells was measured by the flow cytometer. The electro-transfection efficiency was, on average, 50% based on 3 independent experiments with 3 donors (Figure 1a and b). The viability of the plasmid-electroporated cells were always worse than mock-electroporated counterparts. Normalized against mock-electroporated samples, the average frequency of live cells that were electroporated with plasmids was 65% as determined by the live-cell gate on forward versus side scatter (FSC vs SSC) plot by flow cytometer (Figure 1c).

**Figure 1:**
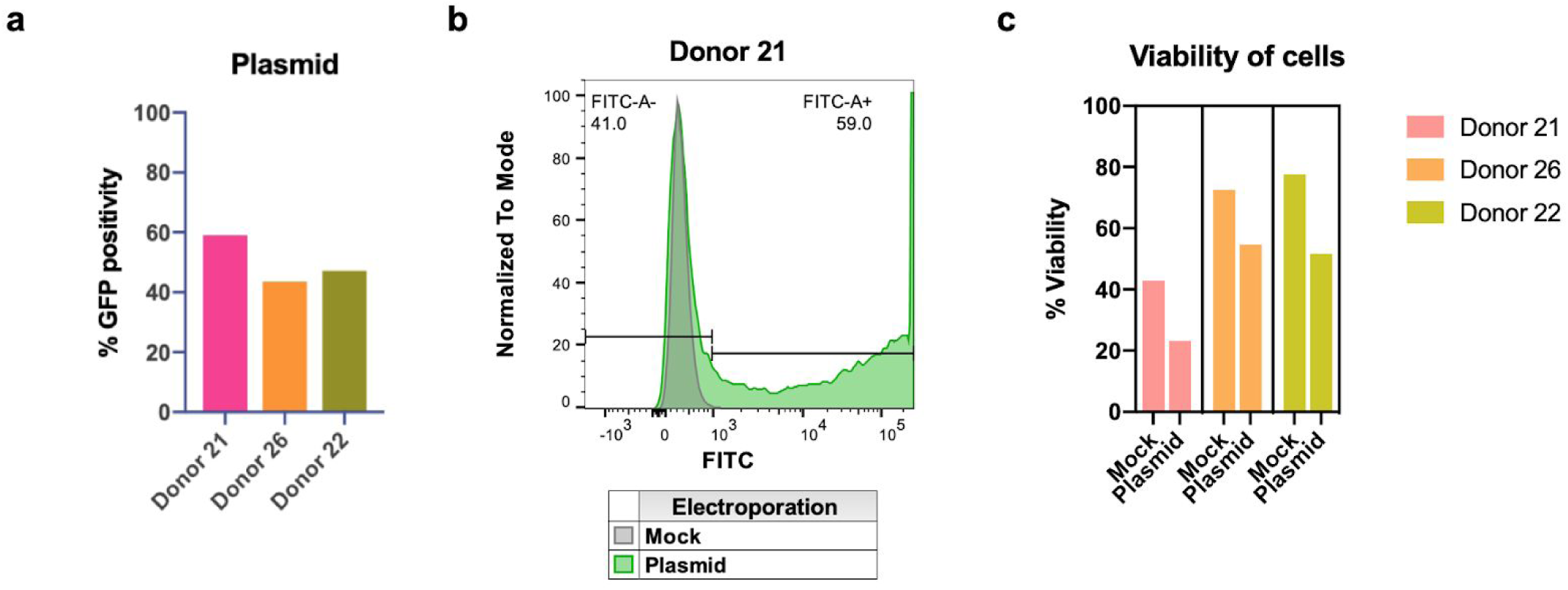
Plasmid electroporation of activated T cells. **a)** Activated T cells were electroporated with a GFP plasmid at a concentration of 7.5 ug DNA per million cells. The frequency of GFP-expressing cells was analyzed by flow cytometer 24 hours after electroporation. The average transfection efficiency was 49.9% **b)** Representative histogram for GFP expression of the plasmid and mock electroporated activated cells. **c)** The viability of plasmid-electroporated cells was consistently lower (on average, 39.5%) than the mock electroporated counterparts (on average, 72%).

#### Plasmid electroporation into unstimulated T cells

To assess the efficiency of electroporation in T cells that have not been activated, we also manipulated unstimulated cells with plasmids. When we kept all of the electroporation settings the same as for activated cells (1600 V 10 ms 3 pulses), there were almost no GFP expressing cells 24 hours after electroporation. These results made us question the electroporation efficiency of unstimulated cells. To better understand the electroporation efficiency of unstimulated cells, we labeled an empty plasmid with Cyanine-5 (Cy5) and electroporated it into both activated and unstimulated cells obtained from the same donors. The frequency of Cy5+ (plasmid positive) cells was higher than 60% for unstimulated cells (Figure 2a and c) and 90% for activated cells (Figure 2b and d) 15 minutes after electroporation. The frequencies of positive cells declined for both groups the next day, but it was still higher than 40% for unstimulated cells (Figure 2a) and almost 80% for activated cells (Figure 2b).

**Figure 2:**
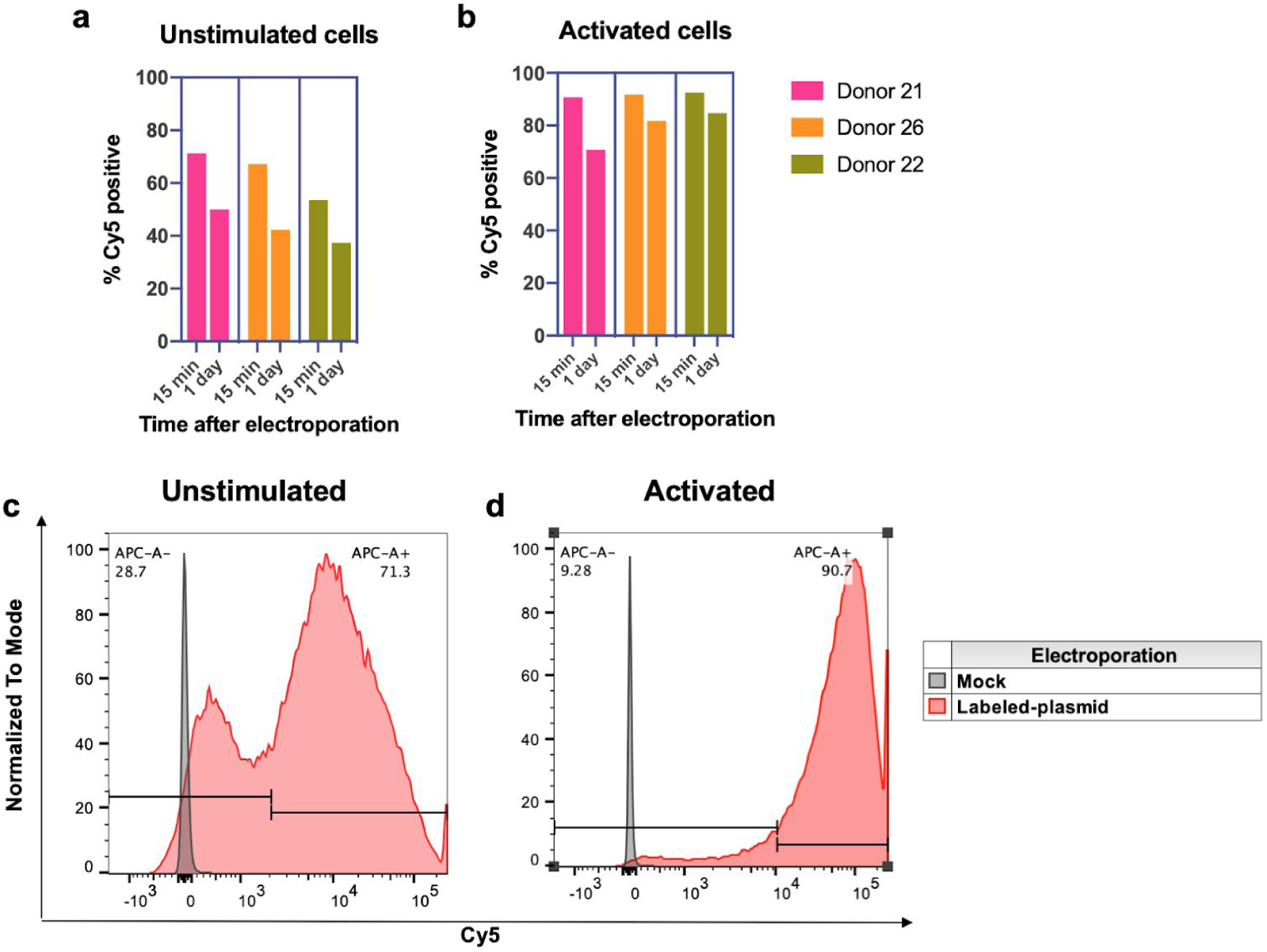
Electroporation of a Cy5-labeled plasmid into activated and unstimulated T cells. Unstimulated **(a,c)** or activated cells **(b,d)** were electroporated with a Cy5-labeled empty plasmid. The frequency of Cy5+ cells was determined by flow cytometer either 15 minutes or 24 hours after electroporation. Both unstimulated (on average, 64%) and activated (on average, 91.6%) cells had a higher frequency of Cy5+ cells on the same day of electroporation, compared to 24 hours after electroporation (43.2% for unstimulated cells and 79% for activated cells). **(c-d)** Examples of electroporation efficiency in unstimulated and activated cells from the same donor 15 minutes after electroporation as detected by flow cytometer.

These results suggested that unstimulated cells were able to take up materials by electroporation but they were not as efficient as their activated counterparts for gene expression. We then imaged the cells by fluorescence microscopy and found that 60% of activated cells were positive for nuclear plasmids whereas unstimulated cells were only 20% positive (Figure S1). Unstimulated cells are smaller compared to activated T cells (Iritani et al., 2002). Therefore, their optimal electroporation settings might be different given that smaller cells require a higher voltage (Shirley et al., 2014; Gehl, 2003). Jay Levy’s group electroporated unstimulated CD8+ T cells with plasmids and achieved 59.6% electro-transfection efficiencies with a viability of 34.6% at 2200 V 20 ms 1 pulse setting using the same electroporation device (Neon, Thermo Fisher) (Liu et al., 2011). When we tried the same settings for electroporating unstimulated cells with our GFP plasmid, we achieved an average of %54.3 electro-transfection efficiency across 3 donors (Figure 3a, orange bars). We also stained the cells for CD45RO and CCR7 surface proteins to estimate the frequency of naive (CCR7+CD45RO-), central memory (CM, CCR7+CD45RO+); effector memory (EM, CCR7-CD45RO+), and effector memory RA (EMRA, CCR7-CD45RO-) subpopulations that were also GFP+ (Sallusto et al., 1999; Mahnke et al., 2013). Our analyses showed that naive cells were mostly GFP positive at this electroporation setting (Figure 3b and d). The viability of plasmid-electroporated cells was around 55%, normalized against mock-electroporated samples (Figure 3c). The viabilities of plasmid-electroporated cells at the 1600V setting were better compared to the ones that were electroporated at the 2200V setting (Figure 3c); however, their electro-transfection efficiency was close to zero (Figure 3a, pink bars).

**Figure 3:**
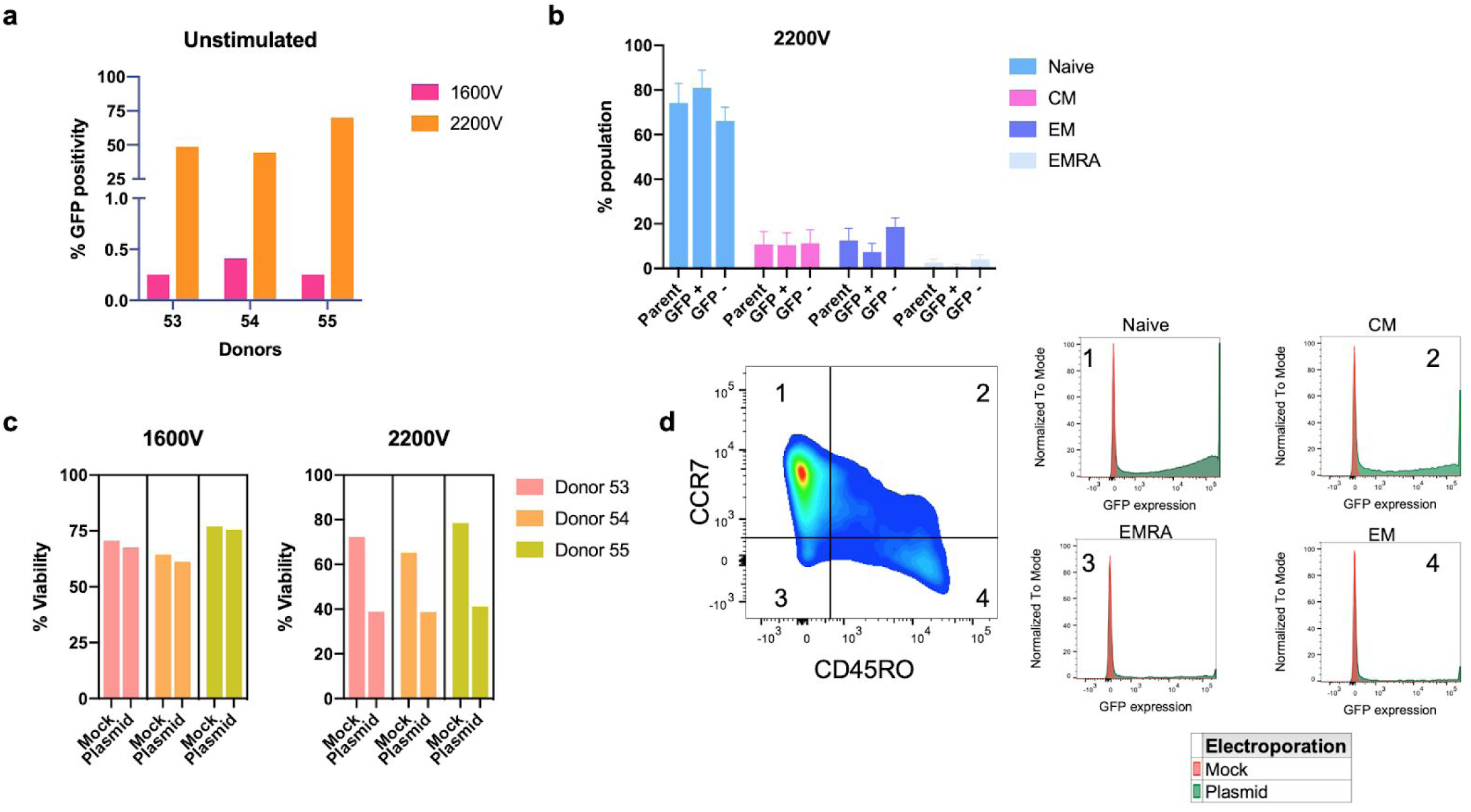
Plasmid electroporation of unstimulated cells at 1600V and 2200V settings. Unstimulated cells from 3 donors were electroporated with a GFP plasmid. The frequency of GFP+ cells were analyzed by flow cytometer 24 hours after electroporation. **a)** Electroporation at 2200V settings is more efficient than the 1600V settings (on average, 54.3% for 2200V and 0.3% for 1600V). **b)** Subpopulations within the unstimulated cell populations were analyzed by staining the cells for CCR7 and CD45RO antibodies. The frequency of GFP+ naive cells was higher than the naive cell frequency in the parent population (80.96% and 74.2%, respectively; n=3). **c)** The viability of the plasmid-electroporated cells was better at the 1600V settings compared to 2200V settings (1600V, mock: 70.6%, plasmid: 68.1%; 2200V mock: 72%, plasmid 39.5%; n=3). **d)** Example of gating strategy with CD45RO and CCR7 staining for estimating the frequency of subpopulations and GFP expression of each subpopulation 24 hours after plasmid electroporation at 2200V.

### Electroporation of mRNA into activated and unstimulated T cells

#### mRNA electroporation into activated cells

Due to decreased viabilities upon plasmid electroporation in both activated and unstimulated T cells, we tried to manipulate the cells with *in vitro* transcribed (IVT) GFP mRNA. We used the same plasmid that we used for plasmid electroporation experiments for in vitro transcribing the mRNA. Activated T cells from 3 donors were electroporated with IVT mRNA (6 ug RNA/million cells) on the second day of activation following debeading. Flow cytometry analysis was then performed on the next day. Using IVT GFP mRNA, we achieved more than 80% GFP+ cells with high viabilities (Figure 4). These results suggested that mRNA electroporation, compared to plasmids, yields better electro-transfection efficiencies and viabilities for activated T cells.

**Figure 4:**
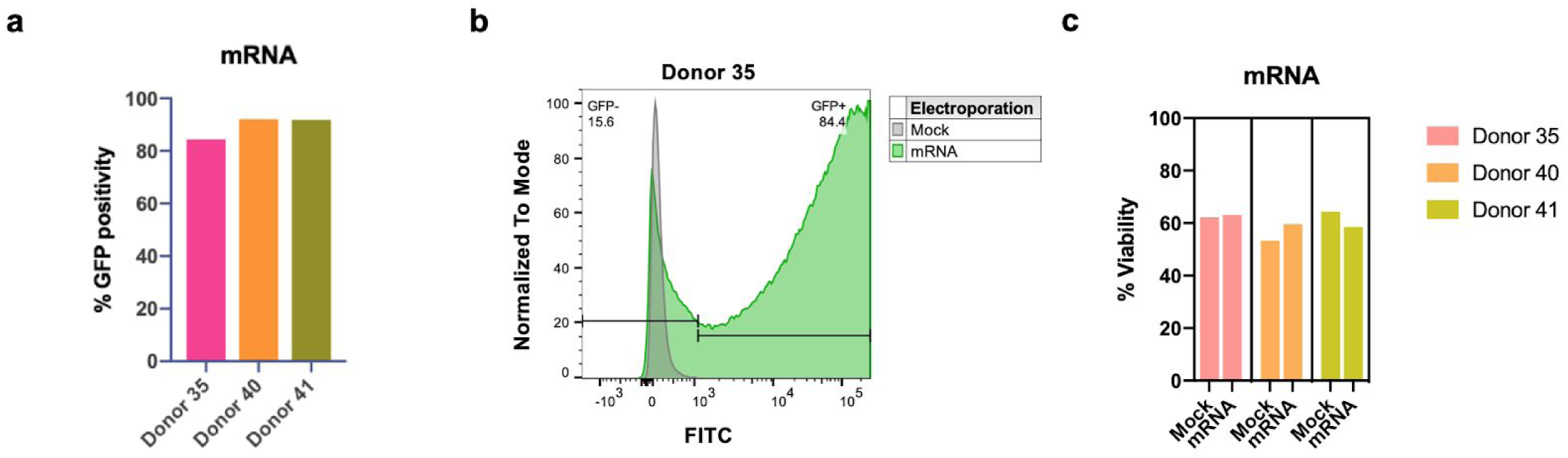
mRNA electroporation of activated T cells. **a)** Activated T cells were electroporated with IVT GFP mRNA (6 ug RNA/million cells) after debeading the cells on the second day of activation. The frequency of GFP+ cells was analyzed by flow cytometer 24h after electroporation and it was higher than 80% across 3 donors. **b)** Representative histogram for electro-transfection efficiency of GFP mRNA electroporated cells. **c)** The viability of the mRNA electroporated cells was similar to the mock-electroporated ones at 24 hours after electroporation.

#### mRNA electroporation into unstimulated cells

Similar to activated cells, manipulating unstimulated cells with plasmids also resulted in decreased viability (Figure 3c, 2200V). mRNA electroporation of activated cells was not as harsh as plasmid electroporation and the frequency of GFP+ cells was also higher (compare 1a and 4a; 1c and 4c). Therefore, we used the same IVT mRNA (8 ug RNA/million cells) to electroporate unstimulated cells to achieve a more efficient and viable electroporation in these cells. We electroporated the cells at both 1600V and 2200V settings to compare the efficiency at two different settings.

mRNA electroporation at 1600V settings resulted in 35% GFP+ cells and no apparent cell death (Figure 5a pink bars and c). 2200V setting resulted in even better electro-transfection efficiencies (>95% GFP+ cells; Figure 5b) and there was still no apparent cell death (Figure 5a orange bars and c). When the subpopulations within the unstimulated cells were analyzed for their GFP expression, we found that the main difference was the GFP-positivity of the naive population between the two electroporation settings (Figure 5d). Our results suggested that 2200V settings were more successful for introducing the mRNA into naive cells.

**Figure 5:**
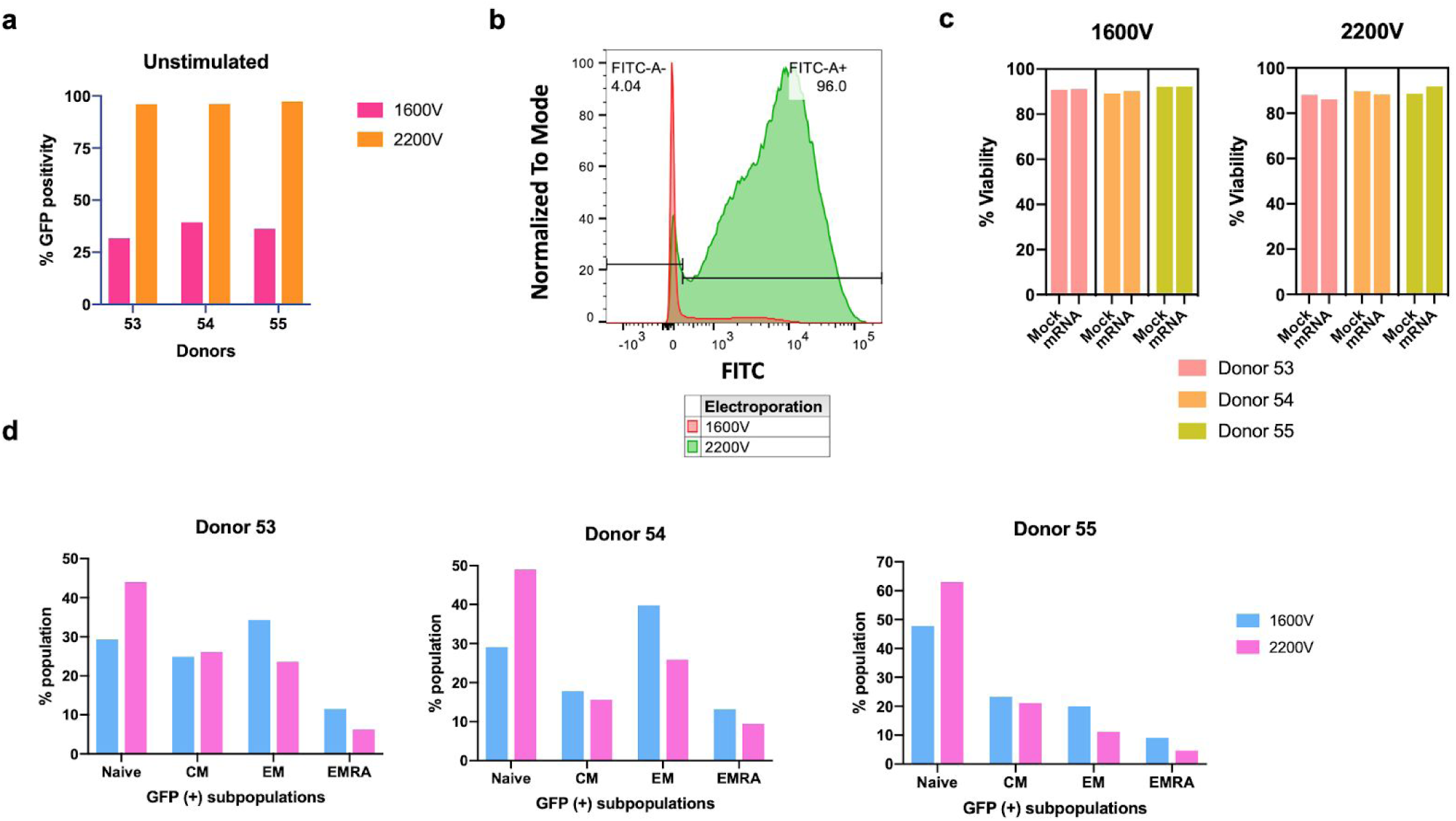
mRNA electroporation of unstimulated cells. Unstimulated T cells were electroporated with IVT GFP mRNA (8 ug RNA/million cells) either at 1600V or 2200V settings. **a)** Electro-transfection efficiency of unstimulated cells was on average 95% with the 2200V settings and 35% with the 1600V settings. **b)** Example of GFP expression 24 hours after electroporation at 1600V or 2200V. The frequencies are shown for the 2200V setting. **c)** Viabilities of the mRNA-electroporated cells were similar to the mock-electroporated controls at both electroporation settings. **d)** Frequencies of naive, CM, EM, and EMRA subpopulations were analyzed based on their GFP positivity after electroporation at 1600V and 2200V. Naive cells were highly GFP+ at 2200V setting. The bar graphs show the frequencies of GFP+ subpopulations in both settings.

### CRISPR in activated and unstimulated T cells

#### Targeting CD4 and CD25 in activated T cells

To accomplish gene editing via CRISPR/Cas9 system in T cells, similar to other cell types, two components should be delivered into the cell: Cas9 and gRNA. These components can be delivered into target cells with viral-vectors or via electroporation. Cas9 and gRNAs can be electroporated as plasmids, as RNA or as Cas9 RNP complex. Among all of these methods, the number of studies using Cas9 RNP in T cells has been increasing for the last couple of years given its efficiency and low toxicity compared to plasmids and also given its transient nature (Schumann et al., 2015; Roth et al., 2018; Seki and Rutz, 2018).

In this study, we also explored the success of gene editing via Cas9 RNP in both activated pan T cells and unstimulated CD4+ T cells. On the second day of CD3/CD28 bead activation, the activated cells from 2 donors were debeaded and electroporated with Cas9 RNPs (7.5 pmol sgRNA and 1250ng Cas9 per 200,000 cells, as recommended by the manufacturer, either against CD25 or CD4). We used one chemically modified synthetic target gene-specific CRISPR RNA (crRNA) per target. Because both target proteins were cell surface proteins, we were able to check the knockout efficiencies by flow cytometer. For each target, we had 3 replicates from both donors (Figure 6a). We achieved a knockout efficiency of 86% for CD4 and 84.4% for CD25 (Figure 6a and b). The cell viabilities were similar to the mock-electroporated samples when checked by flow cytometer 3 days after electroporation (Figure 6c).

**Figure 6:**
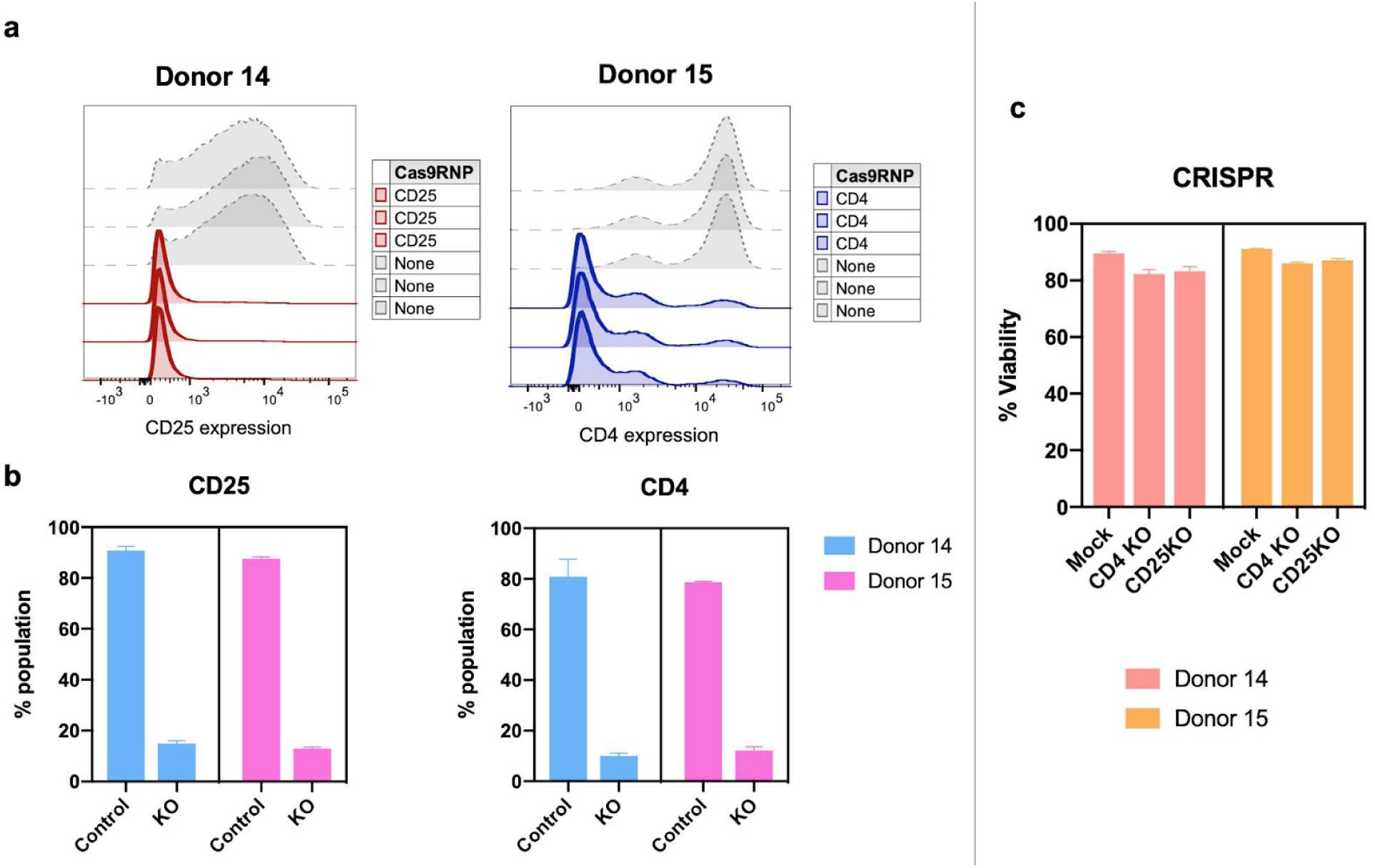
CRISPR in activated T cells. Activated T cells were electroporated with Cas9 RNPs against CD25 or CD4 and the protein levels were measured by flow cytometer 3 days after electroporation. **a)** The efficient knockout of both targets in activated T cells are shown by histograms with three replicates from the same experiment. **b)** CD25+ cell frequencies decreased from 89.1% to 13.9% in CD25-Cas9 RNP electroporated samples and CD4+ cell frequencies decreased from 79.75% to 11.08% in CD4-Cas9 RNP electroporated samples. The bar graphs were plotted using the frequencies of CD25 or CD4 positive cells from each treatment group with 3 experimental replicates and the mean is shown with standard deviation. **c)** Viabilities of the Cas9 RNP-electroporated cells were similar to the mock-electroporated controls at 3 days after electroporation. KO: samples that were electroporated with the corresponding Cas9 RNPs.

#### Targeting CXCR4 and CD127 in unstimulated CD4 (+) cells

To our knowledge, there is only one CRISPR study that showed efficient knockout in unstimulated human T cells (Seki and Rutz, 2018). In the study, the group used Lonza’s Nucleofector to knockout CXCR4, CD127, and CCR7 in human CD4+ T cells by delivering 3 crRNAs per target. They achieved around 90% knockout efficiency with 60% viability, 3 days after electroporation.

To replicate these findings, we used the same crRNA sequences against CXCR4 and CD127. Instead of isolating CD4+ cells directly from fresh PBMCs, we thawed the T cells that we isolated from healthy human blood and then enriched CD4+ cells by depleting CD8+ cells. We then electroporated the unstimulated CD4+ T cells with Cas9 RNPs (3 crRNAs against one gene) using Neon transfection system either at the 1600V or the 2200V setting. The knockout efficiencies were checked by flow cytometer on day 3 and day 6 to account for potentially slow protein turn-over due to the nature of the unstimulated cells. The cells were also stained with CD45RO and CCR7 antibodies to estimate the subpopulation frequencies and the knockout efficiency within each subpopulation.

Using the 1600V settings and CD127 Cas9 RNPs, we did not detect successful knockout events in any of the subpopulations for any of the 4 donors (Figure 7a). However, there was a small decrease in the CD127 protein levels within the CM and EM subpopulations as measured by the mean fluorescence intensity (MFI) using the flow cytometer (Figure 7b). When the same Cas9 RNPs were electroporated into the cells at the 2200V setting, all of the subpopulations–including the naive cells–predominantly lost the CD127 protein at the cell surface (Figures 7c and d).

**Figure 7:**
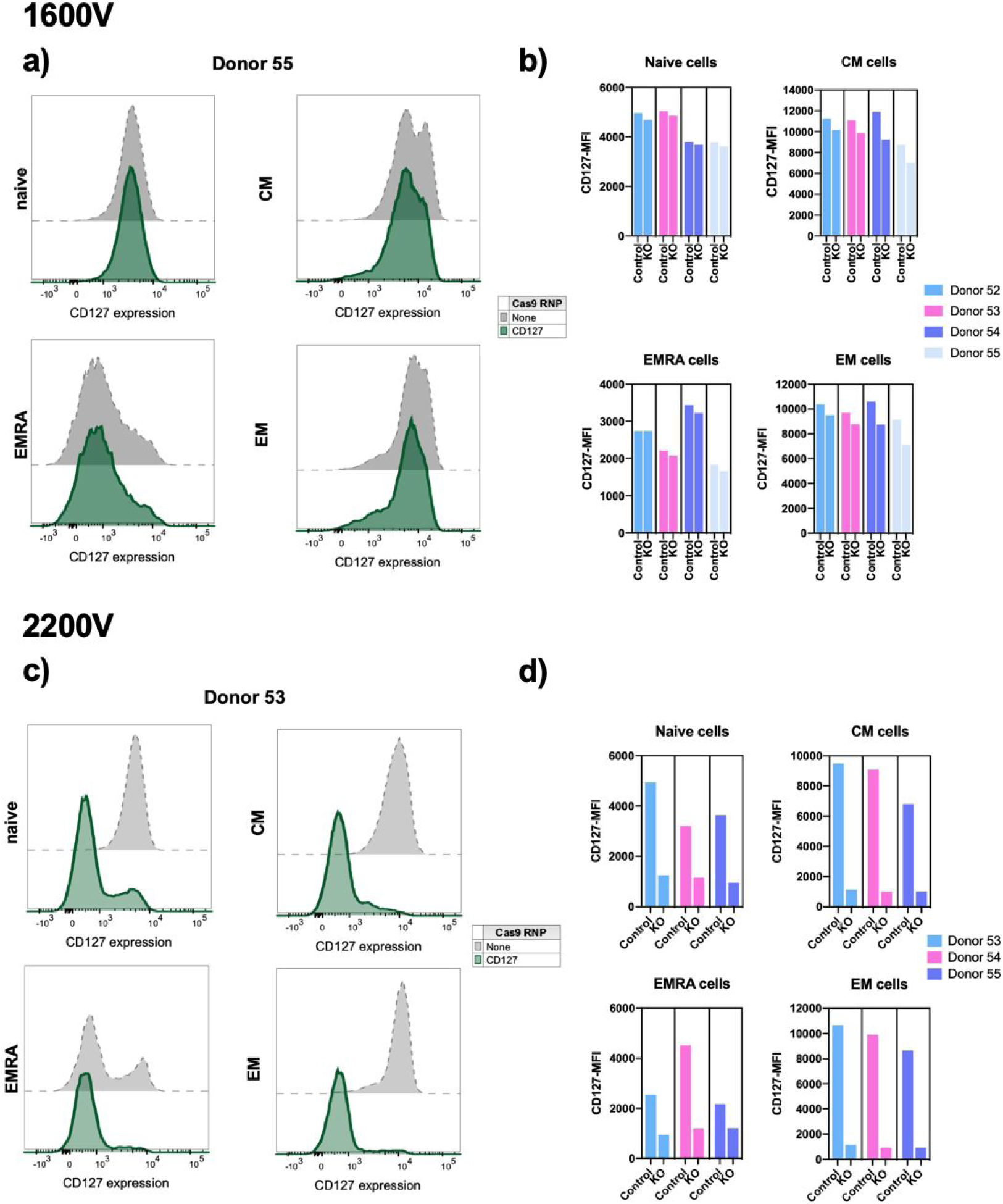
CRISPR in unstimulated T cells: CD127. Unstimulated CD4+ T cells were electroporated with CD127-Cas9 RNPs either at 1600V **(a, b)** or 2200V **(c, d)** setting by Neon electroporation machine. CD127 expression was checked 3 days after electroporation. **(a and b)** For all subpopulations, inefficient CD127 knockout was observed by flow cytometer when Cas9 RNP electroporation was performed at 1600V. (c and d) Successful CD127 knockout was achieved when the same Cas9 RNPs were electroporated into the cells at 2200V. **(b and d)** The bar graphs were plotted using the MFI values of CD127 expression within each subpopulation. Control bars show the MFI values of the mock-electroporated samples.

Similar to the CD127 CRISPR experiments, electroporation at the 1600V setting did not result in efficient knockout of the CXCR4 protein (Figure 8a). Although for some of the donors (*e.g.* donor 54) there was a slight decrease in CXCR4 MFIs within all subpopulations, overall, it was not a successful knockout event when the electroporation was performed at 1600V (Figure 8b).

**Figure 8.**
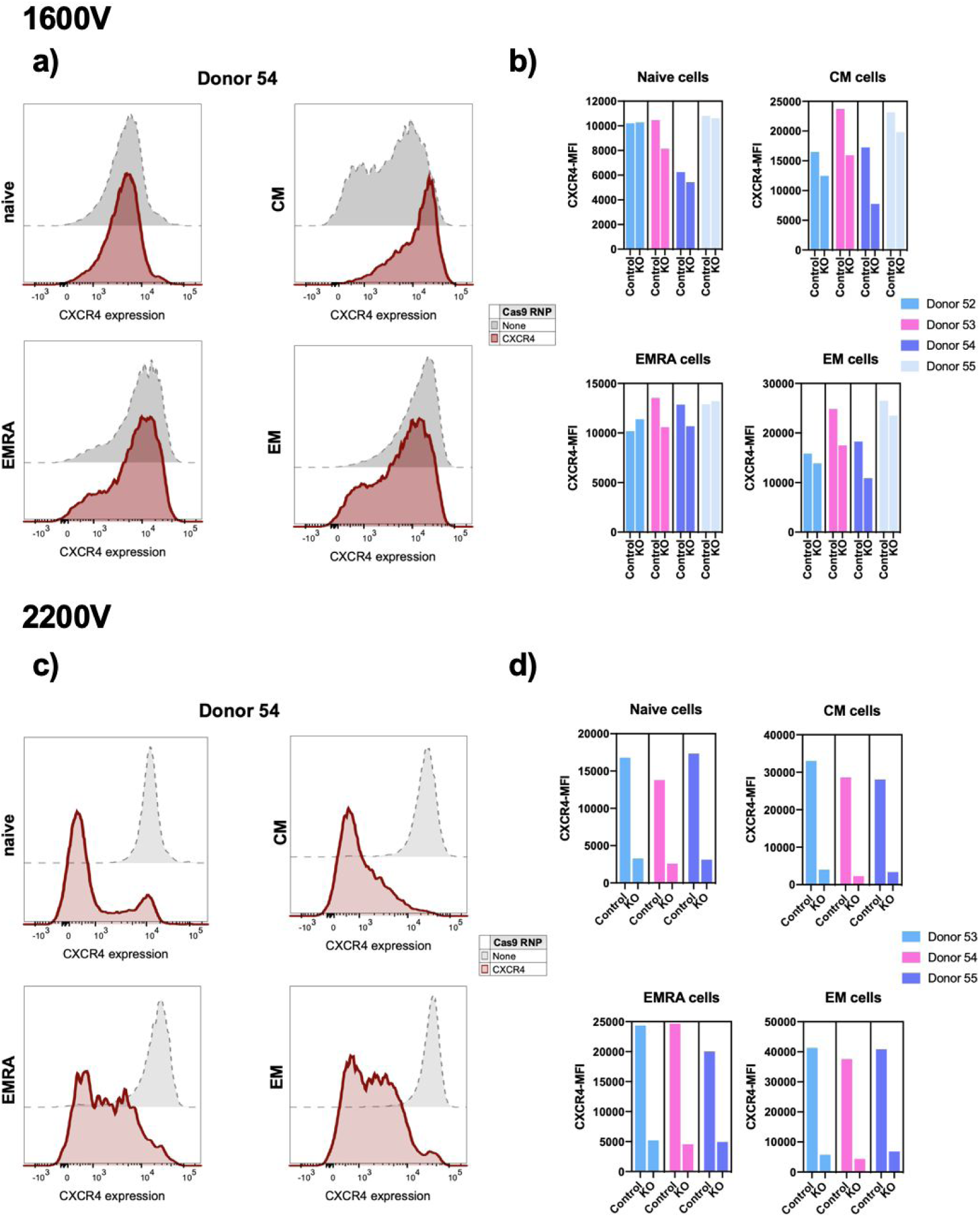
CRISPR in unstimulated T cells: CXCR4. Unstimulated CD4+ T cells were electroporated with CXCR4-Cas9 RNPs either at 1600V **(a, b)** or 2200V **(c, d)** setting by Neon electroporation machine. CXCR4 expression was checked 6 days after electroporation. **(a and b)** For all subpopulations, inefficient CXCR4 knockout was observed by flow cytometer when Cas9 RNP electroporation was performed at 1600V. **(c and d)** Successful CXCR4 knockout was achieved when the same Cas9 RNPs were electroporated into the cells at 2200V. **(b and d)** The bar graphs were plotted using the MFI values of CXCR4 expression within each subpopulation. Control bars show the MFI values of the mock-electroporated samples.

Similar to the CD127 example, electroporating the CXCR4 Cas9 RNPs at 2200V setting resulted in highly efficient knockout of the protein within all subpopulations (Figure 8c and d).

### Using mRNA electroporation for expression of functional TCRs

#### mRNA electroporation optimization of activated mouse T cells

After detecting higher efficiencies and viabilities with mRNA electroporation compared to plasmid electroporation, we decided to electroporate T-cell receptors (TCRs) into human T cells in mRNA format. However, we first wanted to show that this method worked in mouse T cells (isolated from the spleen) with well-characterized mouse TCRs. We started optimizing electroporation settings for activated mouse T cells. For that, we electroporated CD3/CD28-bead-activated mouse CD8+ T cells with GFP mRNA at different voltage settings. Among the five different electroporation settings we have tested, 1300 V 10 ms 3 pulses (1300V) setting gave us the highest cell viability (72.5% viability compared to 77.7% of unelectroporated samples) and the highest efficiency (86.8% FITC positivity) (**Figure 9**).

**Figure 9.**
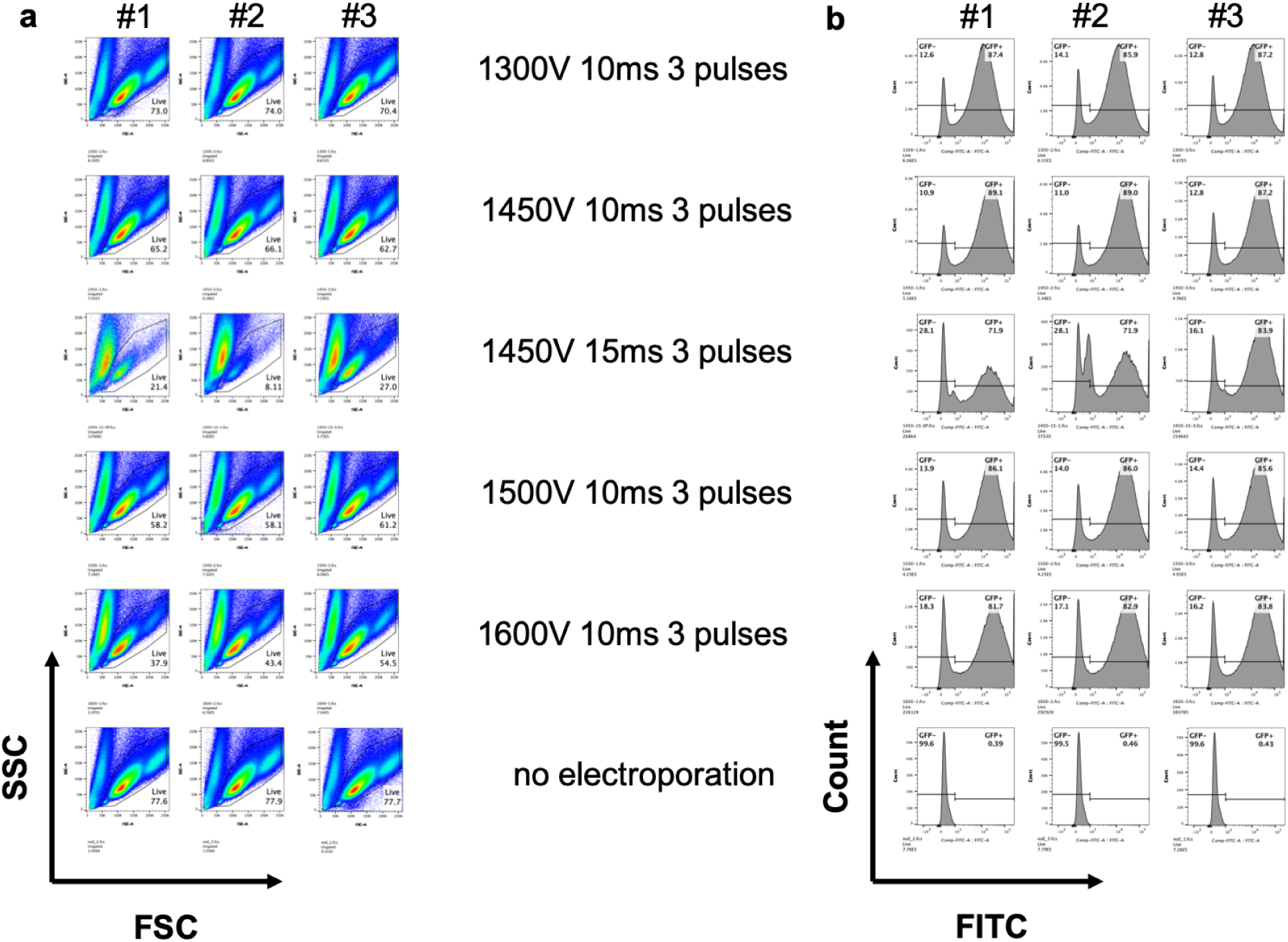
mRNA electroporation optimization of activated mouse CD8+ T cells. Bead-activated mouse CD8+ T cells were electroporated with IVT GFP mRNA (10 ug RNA/million cells) on the fourth day of activation using 5 different electroporation settings. The cells were run on the flow cytometer the next day. **a)** FSC/SSC plots were used to estimate viability. **b)** The electro-transfection efficiency of GFP mRNA electroporated cells was compared based on FITC histograms. The 3 plots next to each electroporation setting show 3 technical replicates.

#### Effect of differently built TCR constructs on expression and function

There are different ways of delivering TCR subunits to cells: (i) alpha and beta subunits can be cloned on individual constructs; (ii) alpha and beta subunits can be cloned on a single construct with a linker sequence in between and alpha subunit can be at the 5’ site; (iii) alpha and beta subunits can be cloned on a single construct with a linker sequence in between and beta subunit can be at the 5’ site. Previously, using retroviral vectors, it has been shown that transducing alpha and beta subunits individually yield higher efficiencies and functionalities in primary murine T cells compared to single vector transduction (Leisegang et al., 2008).

In our study, we picked murine TCR OT-I as our TCR of interest. This MHC class I-restricted TCR recognizes a peptide that is part of the ovalbumin protein (residues 257-264; SIINFEKL). We electroporated OT-I TCR into mouse T cells using the 3 different approaches as described above. We used H2Kb-SIINFEKL multimers to stain SIINFEKL-specific T cells and estimate the fraction of T cells with functional TCRs. We found that electroporating alpha and beta subunit mRNAs separately increased the fraction of cells with functional OT-I TCRs on the cell surface (**Figure 10**). We also found that the beta-linker-alpha construct was superior to alpha-linker-beta construct (20.23% and 0.74% multimer positivity, respectively).

**Figure 10.**
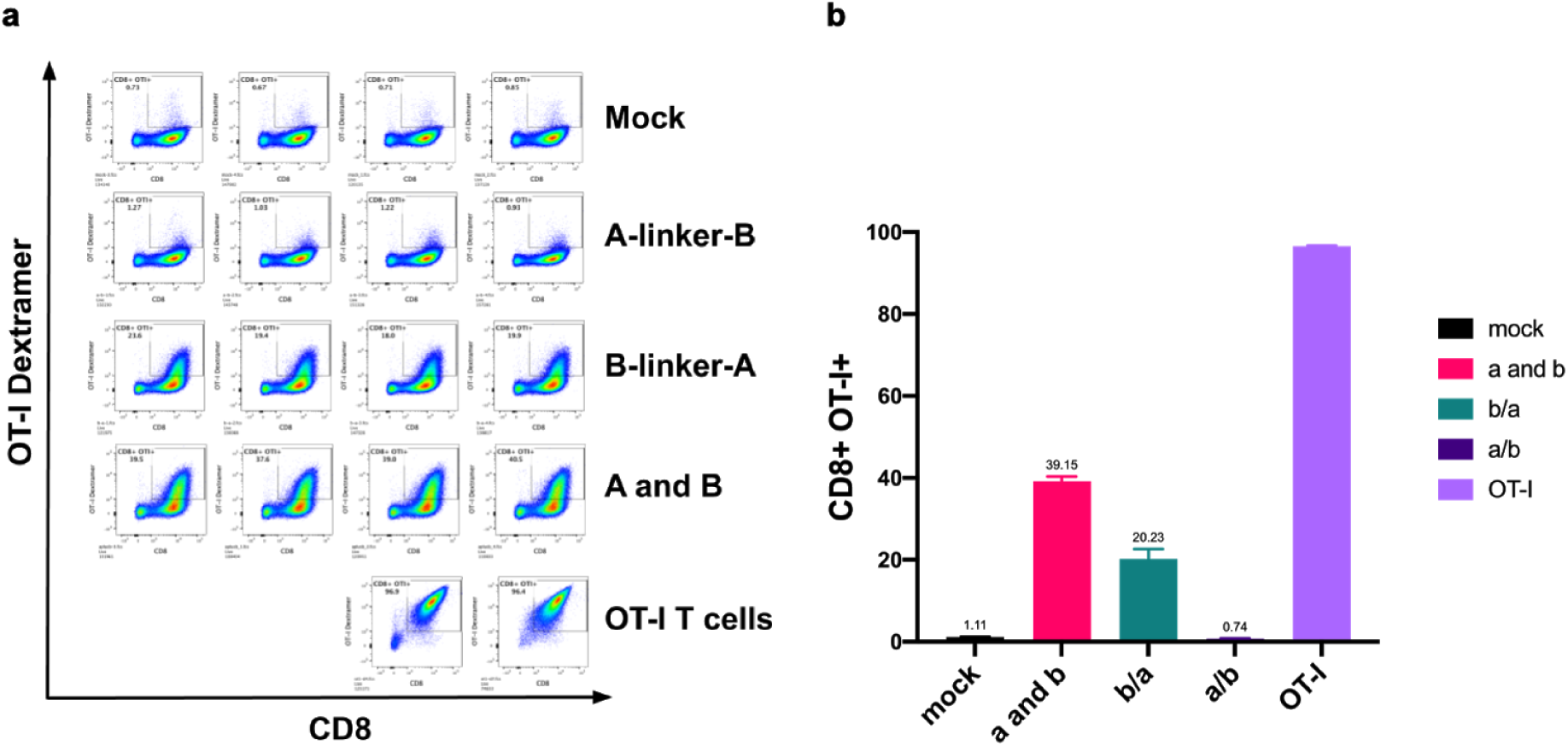
Different constructs of OT-I TCR and their effect on expression and TCR functionality. Bead-activated mouse CD8+ T cells were electroporated with different constructs of OT-I in mRNA form on the fourth day of activation using 1300V setting. **a)** Mock-electroporated cells (negative control), OT-I cells (positive control), and the OT-I TCR electroporated cells were stained with CD8 and H2Kb-SIINFEKL dextramers to estimate the frequency of functional OT-I TCRs on the cell surface. **b)** The data was plotted on a bar graph to summarize the results. n=4 for all samples except OT-I T cells. **A-linker-B**: OT-I TCR alpha-p2a-OT-I TCR beta; **B-linker-A**: OT-I TCR beta-p2a-OT-I TCR alpha; **A and B**: OT-I TCR alpha and OT-I TCR beta mRNAs electroporated individually.

### Cytotoxicity assays with TCR-electroporated T cells and peptide-pulsed cancer cells

#### Optimization of the cytotoxicity assay with OT-I mouse T cells

Our goal was to assess whether OT-I TCR electroporated mouse T cells were cytotoxic against SIINFEKL-presenting cancer cells. First, we wanted to make sure our cytotoxicity assay was capable of detecting T cell-mediated killing when we co-cultured T cells from OT-I transgenic mice and pulsed the target cancer cells with SIINFEKL peptide. For the cancer cell line side, we used the murine colon adenocarcinoma cell line, MC38, after evaluating its H2Kb expression and SIINFEKL presentation (**Figure S2**). Because our cytotoxicity assay was based on LDH release and LDH had a half-life of 9 hours, we set two-time points for the co-culture experiment to be able to capture the killing. We found that 6-hour co-culturing was able to capture the cytotoxicity activity better compared to the overnight co-culturing. (**Figure S3**).

#### Cytotoxicity assays with OT-I TCR electroporated mouse T cells

Our test experiment using the H2Kb-SIINFEKL multimer staining suggested that the best way to introduce OT-I TCR was by electroporating alpha and beta subunits individually. Therefore, for the co-culture experiments, we followed this strategy and also included T cells from OT-I transgenic mice as a positive (SIINFEKL-presentation-reactive) control group. For the wild type T cell electroporation, we used splenocytes from C57BL/6J (WT) mice and activated them with CD3/CD28 beads for at least 2 days. CD8+ T cells were enriched after debeading and mRNA electroporation was done 5 days after activation. We started the co-culture the next day of mRNA electroporation and, on the same day, we also stained SIINFEKL-specific T cells with the multimer to see our electroporation efficiency (**Figure 11a**). Similar to our previous results, the multimer staining of the OT-I TCR electroporated cells was around 40%. We compared cytotoxicities of electroporated T cells, T cells from OT-I mice, and the WT T cells against SIINFEKL-pulsed MC38s in different T cell to MC38 ratios. Although H2Kb-SIINFEKL multimer staining of electroporated T cells and T cells from OT-I mice was different (**Figure 11a**), the cytotoxicity assay results suggested that these two groups were similar in their functionality (**Figure 11b**).

**Figure 11.**
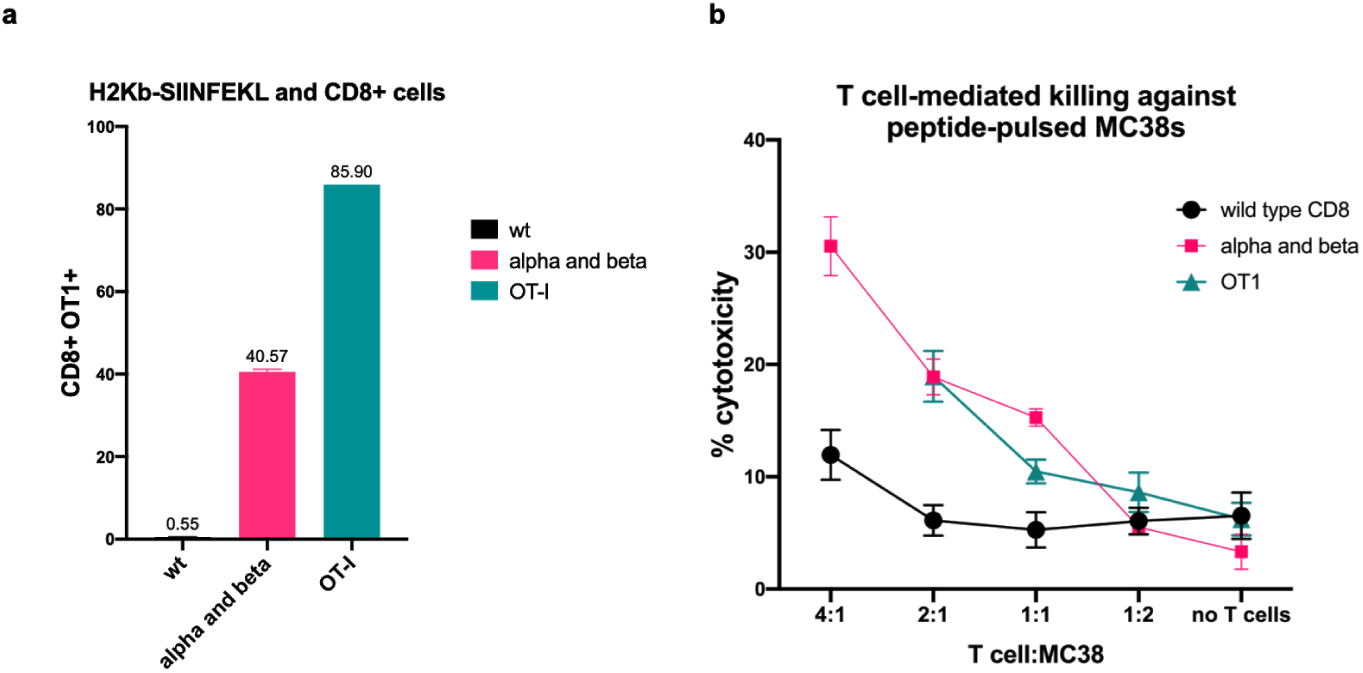
Cytotoxicity assays with OT-I TCR electroporated mouse CD8+ T cells and peptide-pulsed MC38s. Bead-activated CD8+ T cells were co-electroporated with OT-I TCR alpha mRNA and OT-I TCR beta mRNA (10 ug total RNA/million cells). **a)** The next day, cells were stained with H2Kb-SIINFEKL dextramer and CD8 antibody to estimate the frequency of functional OT-I TCRs on the cell surface by flow cytometer. OT-I cells served as the positive control group. wt: unelectroporated mouse CD8+ cells. n=3 for wt and alpha and beta. **b)** Cytotoxicity assay was set up with different ratios of T cells:MC38s and the co-culture were kept for 8 hours. MC38s were pulsed with 10 uM SIINFEKL peptide and the pulsing and T cell addition was performed simultaneously. Percent cytotoxicity was estimated based on LDH release using a plate reader. n=6.

#### Cytotoxicity assays with OT-I TCR electroporated human T cells

##### Estimating cytotoxicity by LDH-release assay

The data from the mouse T cell experiments suggested that OT-I TCR could be electroporated into activated WT mouse CD8+ T cells in mRNA form and these electroporated WT cells could kill the target cells as efficiently as CD8+ T cells from OT-I transgenic mice. Building upon this information, we wanted to repeat the same experiments using human CD8+ T cells. The first thing we checked was the availability of functional mouse TCR (OT-I) on human CD8+ T cell surface upon electroporation. We isolated pan T cells from healthy human donor blood and activated these cells with anti-CD3/CD28 beads. After debeading, we enriched for CD8+ T cells and electroporated them with individual OT-I alpha and beta mRNAs simultaneously. Using the H2Kb-SIINFEKL multimer, we showed that SIINFEKL-specific CD8+ T cell frequencies were similar across 4 donors (**Figure S4**).

Multimer-staining data suggested that human T cells could be electroporated with murine OT-I TCR and these TCR were functional. To test this, we set up co-cultures of OT-I TCR electroporated human CD8+ T cells and SIINFEKL pulsed or unpulsed MC38s. When T cells are isolated from buffy coats as pan T cells, most of them are CD4+ T cells (Aksoy et al., 2018). Therefore, with our experiments having enough CD8+ T cells to start the co-culture was generally a problem. With our first experiment, we had enough OT-I TCR electroporated human CD8+ T cells to set up a co-culture experiment with 4:1 T cell to MC38 ratios. We checked the electroporation efficiency by staining the electroporated cells with the H2Kb-SIINFEKL multimer and detected 59% multimer positivity (**Figure 12a**). These electroporated CD8+ T cells specifically killed the SIINFEKL-pulsed MC38s in 8 hours and the efficiency was better when T cell to MC38 ratios was higher (**Figure 12b**). When we repeated the experiment with another donor, we did not have enough CD8+ T cells to reach 4:1 T cell to MC38 ratio, we could only reach to 2.6:1. multimer staining of the electroporated cells showed 45% multimer positivity (**Figure 12c**). Similar to our first experiment, within all of the co-culture ratios, electroporated CD8+ T cells killed the pulsed MC38 more than the unpulsed ones (**Figure 12d**). Our third repeat with the same setup and with different donor’s CD8+ T cells yielded 33% multimer positivity (**Figure 12e**). With this experiment, we had enough cells to set up the co-culture with 4:1 T cell to MC38 ratios and we could compare the cytotoxicity levels from an 8-hour co-culture to an overnight co-culture. To our surprise, OT-I electroporated CD8+ T cells did not kill the pulsed MC38s more than unpulsed MC38s at ratios other than 2:1 (**Figure 12f**). The specific killing measurements benefited from the overnight co-culture: at all T cell to MC38 ratios, OT-I electroporated CD8+ T cells killed the pulsed cells more than the unpulsed ones (**Figure 12g**).

**Figure 12.**
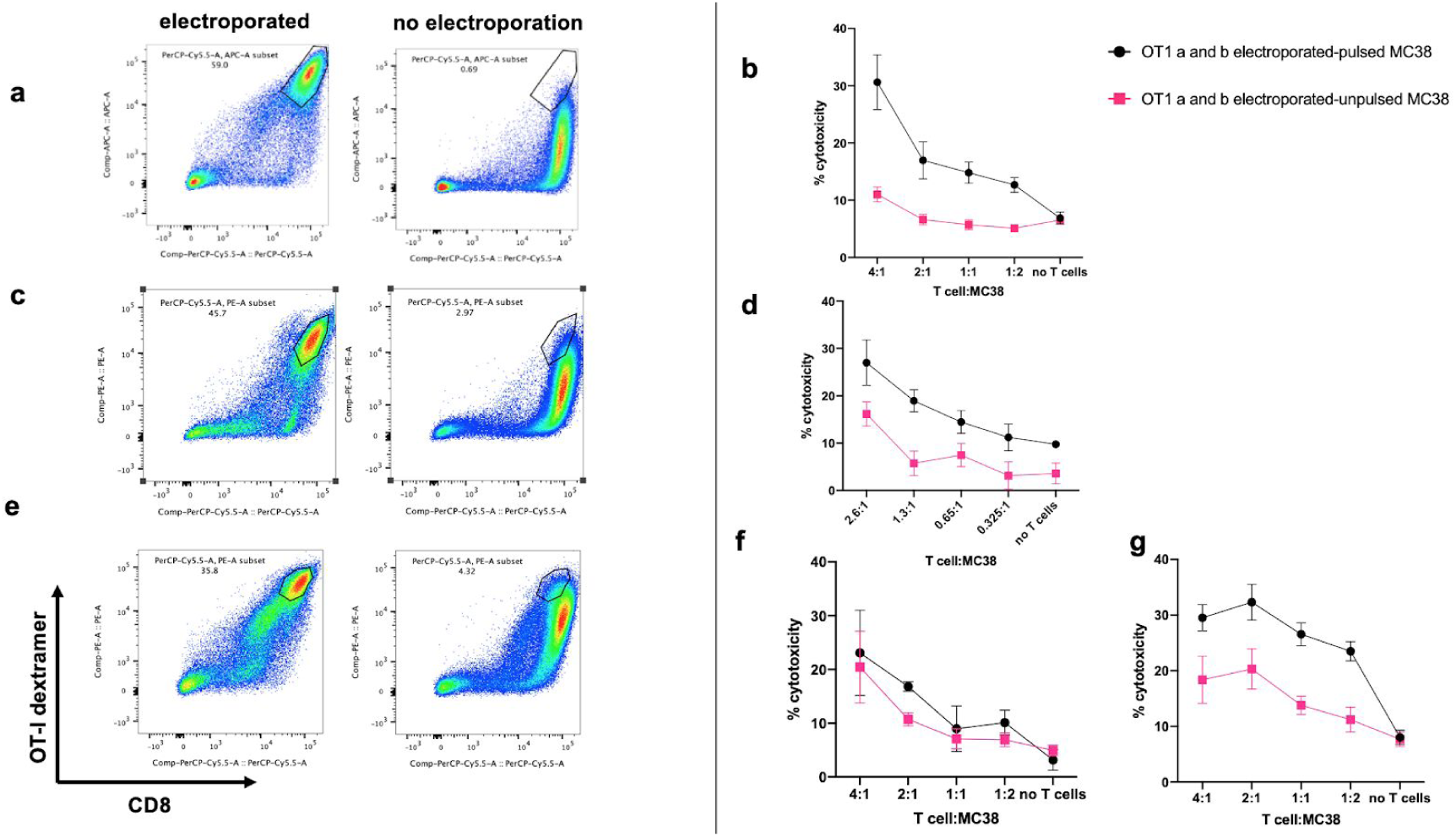
Cytotoxicity assays (LDH release) with OT-I TCR electroporated human CD8+ T cells and peptide-pulsed or unpulsed MC38s. Bead-activated human CD8+ T cells from 3 donors were co-electroporated with OT-I alpha and beta mRNAs (10 ug total RNA/million cells). **a,c,e)** The next day, electroporated and unelectroporated cells were stained with H2Kb-SIINFEKL dextramer and CD8 antibody to estimate the frequency of functional OT-I TCRs on cell surface by flow cytometer. **b,d,f)** Cytotoxicity assays were set up with different ratios of T cells:MC38s and the co-culture were kept for 8 hours and overnight for **g.** For the pulsed MC38 group, the cells were pulsed with 10 uM SIINFEKL peptide and the pulsing and T cell addition was performed simultaneously. Percent cytotoxicity was estimated based on LDH release using a plate reader. n=6.

#### Estimating cytotoxicity by flow cytometer

The cytotoxicity assay results suggested that OT-I TCR electroporated human CD8+ T cells could kill the target cells with varying efficiency and specificity from donor to donor. Because having enough CD8+ T cells was the biggest hurdle for this cytotoxicity assay, we decided to approach the same question with a different technique and solution. Instead of co-culturing increasing ratios of T cells:MC38s, we decided to select the highest ratio (8:1) and only use that one for our co-culture experiment.

For this trial, we isolated pan T cells from 3 donors and activated all of them at the same time. On the second day of activation, we debeaded the cells and enriched for CD8+ T cells. Following the enrichment, we co-electroporated these cells with OT-I TCR alpha and beta mRNAs (10 ug total RNA/million cells). We also included activated mouse CD8+ T cells and electroporated them with OT-I TCR alpha and beta mRNAs and also prepared activated T cells from OT-I transgenic mice to serve as our positive control group for cytotoxicity. The next day, we co-cultured electroporated CD8+ T cells with MC38s at a T cell:MC38 ratio of 8:1. For each group, we kept one set of MC38s unpulsed, and pulsed the other set with SIINFEKL peptide simultaneously with T cell addition. After incubating the co-cultures overnight, we collected all of the cells both by pipetting and trypsinization. We stained the cells with CD3 and CD8 cell surface antibodies to be able to distinguish between the MC38 cells (target: CD3- CD8-) and T cells (effector: CD3+ CD8+). By using the frequency of both live and CD3- CD8- cells, we calculated an estimated number for MC38 cell counts under each co-culture group. Unpulsed MC38 cell counts were ∼110,000 when they were co-cultured with OT-I TCR electroporated human T cells, whereas the cell count decreased to 24,000 when MC38s were pulsed indicating specific killing (Figure 13a). We, then, normalized the pulsed MC38 counts by the unpulsed MC38 counts for each donor. The average cytotoxicity was calculated as 78.8% for 3 donors (Figure 13b).

**Figure 13.**
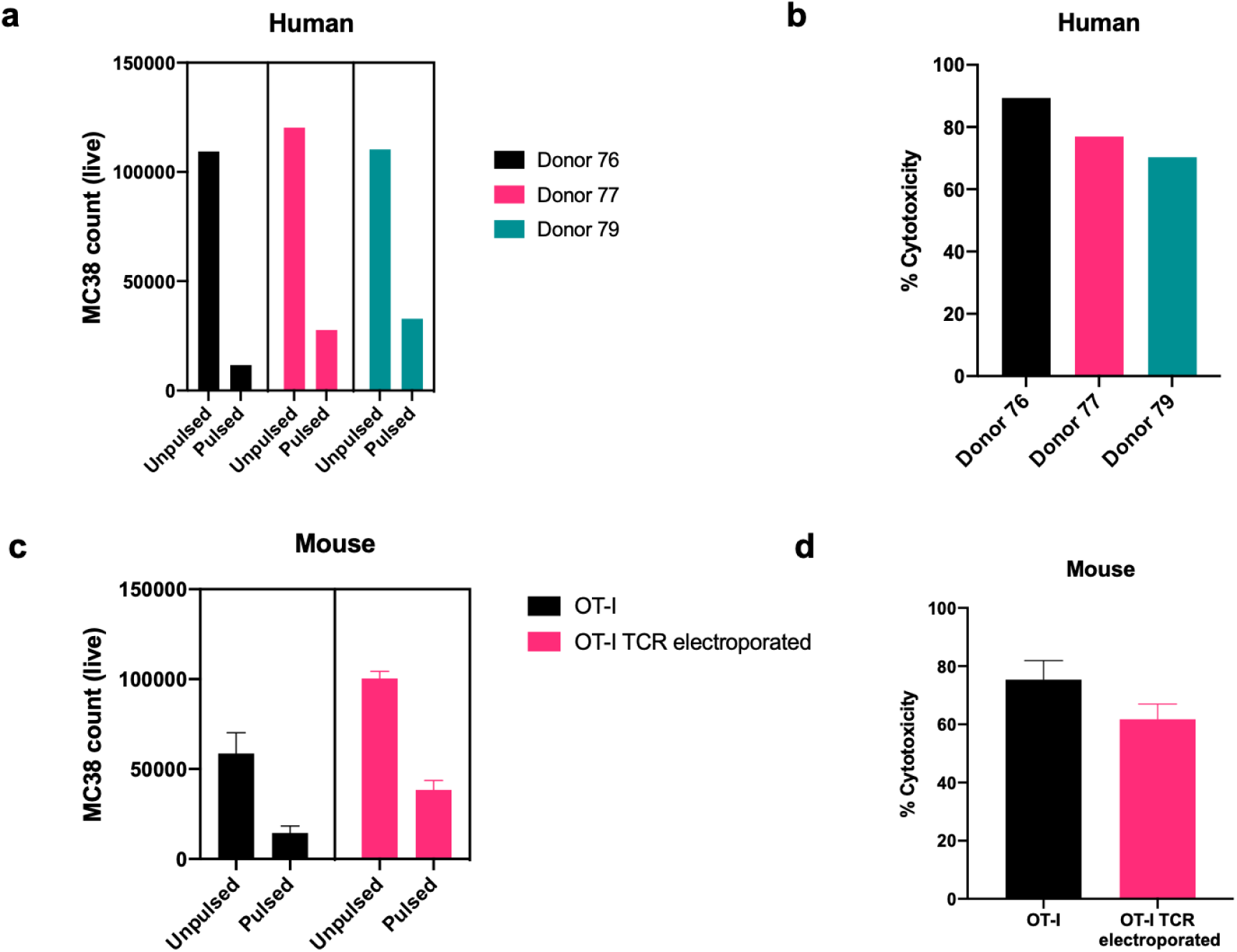
Cytotoxicity assays (flow cytometer) with OT-I TCR electroporated human/mice CD8+ T cells and peptide-pulsed or unpulsed MC38s. Bead-activated human and mouse CD8+ T cells were co-electroporated with OT-I alpha and beta mRNAs (10 ug total RNA/million cells). The next day, electroporated cells and CD8+ cells from OT-I transgenic mice were co-cultured with SIINFEKL-pulsed or unpulsed MC38s with a ratio of 8:1. Following overnight incubation, cells were analyzed by flow cytometry and MC38 cell counts were estimated based on anti-CD3/CD8 double negative staining. **a)** Pulsed and unpulsed MC38 cell counts after co-culturing with OT-I TCR electroporated human T cells. **b)** Estimated cytotoxicity of each donor T cells against pulsed MC38s. **c)** Pulsed and unpulsed MC38 cell counts after co-culturing with transgenically OT-I TCR-harboring CD8+ T cells or with OT-I TCR electroporated wt CD8+ cells. **d)** Estimated cytotoxicity of OT-I TCR-harboring CD8+ T cells and OT-I TCR electroporated wt CD8+ cells (n=3 for c and d).

When we co-cultured MC38 with mouse CD8+ T cells from OT-I transgenic mice, the average unpulsed MC38 cell count was 60,000 however, the cell count decreased to 14,000 when the MC38s were pulsed (Figure 13c, black bars). Similarly, the average unpulsed MC38 cell count was 100,000 when the cells were co-cultured with OT-I TCR electroporated mouse CD8+ T cells and the cell count decreased to 38,000 when the MC38s were pulsed with the SIINFEKL peptide (Figure 13c, pink bars). We, then, normalized the average pulsed MC38 counts by the average unpulsed MC38 counts for each group. The cytotoxicities of CD8+ cells were calculated as 75% and 61% for OT-I transgenic mice and OT-I TCR electroporated WT ones, respectively (Figure 13d).

## Discussion

Achieving successful genetic manipulation of primary human T cells is of importance to both basic immunology research and clinical applications involving genetically-altered human T cells. In this study, we characterized the most efficient and less cytotoxic ways of electroporating unstimulated and CD3/CD28 bead-activated T cells. By using our electroporation device (Neon, Thermo Fisher) at two different electroporation settings, 1600 V 10 ms 3 pulses (1600V) for activated cells and 2200 V 20 ms 1 pulse (2200V) for unstimulated T cells, we achieved high electro-transfection efficiencies through delivering in vitro transcribed (IVT) mRNA or synthetic Cas9, to both activated and unstimulated cells. Plasmid electroporation yield (50-55%) was relatively low compared to these two methods for both unstimulated and activated T cells. We also optimized the mRNA electroporation for activated primary mouse CD8+ T cells and used that setting 1300V 10 ms 3 pulses (1300V) for electroporating TCRs into mouse T cells.

Our first attempt of plasmid electroporation in unstimulated cells failed when the electroporation was performed at the 1600V settings. Observing almost 0% efficiency upon plasmid electroporation made us question the abilities of unstimulated cells to take up material by electroporation. We then electroporated activated and unstimulated cells with a fluorescently-labeled empty plasmid. The flow cytometry results showed that unstimulated cells were, indeed, able to take up the labeled plasmid at a level similar to activated cells (Figure 2). Then, we repeated the experiment and imaged the cells 24 hours after electroporation. Imaging results showed that 60% of the activated cells had plasmids in their nucleus whereas the frequency was only 20% for the unstimulated cells (Figure S1). Since unstimulated cells are, on average, smaller than the activated cells, we wanted to test whether a higher voltage setting would improve the efficiency as others have noted on cell size and electroporation efficiency (Shirley et al., 2014; Gehl, 2003). We then performed electroporation at 2200V as was suggested by Jay Levy’s group for plasmid electroporation of unstimulated CD8+ T cells using the Neon electroporation machine (Liu et al., 2011). By electroporating unstimulated cells at the 2200V setting, we achieved a relatively higher efficiency even within the naive subpopulation (overall efficiency: 54.3% versus 0.3% at 2200V versus 1600V, respectively). These results suggested that at 2200V setting more plasmids were introduced into the cells and, in return, more plasmids localized to the nucleus. Similarly, we were able to get relatively high Cas9 RNP-mediated KO in naive cells with the 2200V setting. Using CD4+ unstimulated human T cells, we were able to knockout CXCR4 and CD127 genes in both naive cells and memory cells with similar efficiencies (Figures 7 and 8).

By using our optimized mRNA electroporation settings and OT-I as our TCR of interest, we also showed that functional TCRs can be electroporated into both human and mouse CD8+ T cells. We generated different constructs of the OT-I TCR based on Leisegang and others’ work and electroporated these different constructs in mRNA form into activated mouse CD8+ T cells (Leisegang et al., 2008). Out of the 3 different delivery methods of the TCR mRNA (alpha and beta separate; alpha and beta on a single construct and alpha is at 5’ position; alpha and beta on a single construct and beta is at 5’ position), co-electroporating alpha and beta subunits of the OT-I TCR gave us the best multimer-staining with H2Kb-SIINFEKL multimers (Figure 10). Although multimer staining indicates the presence of a functional TCR pair on the cell surface, we also set up cytotoxicity assays with these OT-I TCR electroporated cells to show that the expression of these functional TCRs actually lead to T-cell-mediated cytotoxicity for the SINFEKL-presenting cells. We first tested the cytotoxicity of WT mouse CD8+ T cells that were electroporated with the OT-I TCR and compared their cytotoxicity to the OT-I CD8+ T cells. The cytotoxicities of the T cells were tested against SIINFEKL-pulsed MC38s. Although their multimer-staining efficiencies were different (Figure 11a), these two different groups of cells (electroporated WT cells and cells from OT-I mice) killed the target cells with similar cytotoxicities (Figure 11b). We then tried the same setup using human T cells. By isolating T cells from different donors and by using the H2Kb-SIINFEKL multimer staining, we showed that functional mouse TCRs could be introduced onto human CD8+ T cells by mRNA electroporation. We setup cytotoxicity assays with OT-I TCR electroporated human CD8+ T cells in two different ways: Our first setup relied on LDH-release assay from the dead cells by a plate reader and the second assay relied on the total cell count that was left in the well after an overnight co-culture incubation by flow cytometry. The issue with the first method was the half-life of the LDH (9 hours) and the different cytotoxic kinetics of different donor’s T cells. With some of the donors, an 8-hour co-culture incubation was sufficient whereas with some overnight incubation was necessary (Figure 12). We then switched to the flow cytometry-based method in which there was no bias against slow-killers or fast-killers. Using T cells from 3 different donors, we showed that all these donor T cells armed with mouse OT-I TCRs could kill the cognate antigen-presenting target cells with similar cytotoxicities after overnight co-culturing (Figure 13). The ability to express functional TCRs at high efficiencies through mRNA electroporation without the need for knocking out the endogenous TCR can facilitate testing candidate TCRs for their reactivity against arbitrary targets.

Electroporation-based transfection of primary cells has been around for decades but its utility as a non-viral alternative to genetic manipulation of human primary T cells has recently been re-evaluated. This is mostly due to the emergence of highly efficient CRISPR/Cas9-mediated gene knockout techniques and their potential for studying basic T cell biology and translational application for T-cell-mediated immunotherapies. Although many other groups have attempted to show the utility of electro-transfection in (mostly activated) human primary T cells, the use of this technique has not been extensively characterized in unstimulated T cells side-by-side with the activated ones. In this study, we systematically profiled the genetic manipulation efficiency of unstimulated and activated T cells through electro-transfection to better evaluate their utility for basic T cell biology and its feasible translation for clinical applications. We show that both electroporation of IVT mRNA for transient gene expression and Cas9 RNP for gene knockout are highly efficient not only in the activated but also in unstimulated cells, including naive T cells. We expect to see wide adoption of these techniques in the near future.

## Materials and Methods

### Cells and cell lines

#### Human primary T cells

PBMCs were isolated from healthy human donors (purchased from Plasma Consultants LLC, Monroe Township, NJ) by Ficoll centrifugation (Lymphocyte separation medium; Corning, Corning, NY). T cells were isolated using Dynabeads Untouched Human T Cells Kit (Thermo Fisher, Waltham, MA) or by using StemCell’s EasySep™ Human T Cell Isolation Kit (Vancouver, Canada) using the manufacturer’s protocols. Isolated T cells were kept in T cell media: RPMI with L-glutamine (Corning), 10% fetal bovine serum (Atlas Biologicals, Fort Collins, CO), 715 uM 2-mercaptoethanol (EMD Millipore), 25 mM HEPES (HyClone, GE Healthcare, Chicago, IL), 1% Penicillin-Streptomycin (Thermo Fisher), 1X sodium pyruvate (HyClone, GE Healthcare, Chicago, IL), and 1X non-essential amino acids (HyClone, GE Healthcare). T cells were activated for 2 days with anti-CD3/CD28 magnetic dynabeads (Thermo Fisher) at beads to cells concentration of 1:1, with a supplement of 200 IU/ml of IL-2 (NCI preclinical repository).

Protocol details:

- Culture media: <u>DOI:10.17504/protocols.io.qu5dwy6</u>
- PBMC isolation from buffy coat: <u>DOI:10.17504/protocols.io.qu2dwye</u>

#### Mouse T cells

Mouse splenocytes were a gift from Rubinstein lab (WT: C57BL/6J, Jackson lab, #000664) and Paulos lab (OT-I transgenic mice: C57BL/6-Tg(TcraTcrb)1100Mjb/J, Jackson lab, #003831). WT splenocytes were cultured at 1 million cells per ml and activated with anti-CD3/CD28 magnetic dynabeads at beads to cells concentration of 1:1, with a supplement of 200 IU/ml of IL-2. Splenocytes from OT-I transgenic mice were activated with the SIINFEKL peptide at 10uM concentration. Bead-activated cells were debeaded on day 3, and all of the activated cells were supplemented with 200 IU/ml of IL-2 on a daily basis starting from day 3. The cell concentration was adjusted to be 500,000-1,000,000 cells per ml.

#### MC38 cell line

MC38 mouse colon adenocarcinoma cells were a gift from the Rubinstein lab at the Medical University of South Carolina. The cells were grown in Dulbecco’s Modified Eagle Medium (DMEM) with high glucose (4500 mg/L) and L-glutamine (4.0 mM) and the media was supplemented with 10% fetal bovine serum (Atlas Biologicals) and 1% Penicillin-Streptomycin (Thermo Fisher).

### Plasmids

- pcDNA3.3_NDG was a gift from Derrick Rossi (Addgene plasmid #26820) (Warren et al., 2010).
- pCMV6-Entry Tagged Cloning Vector was purchased from OriGene (#PS100001).
- murine TCR OTI-2A.pMIG II was a gift from Dario Vignali (Addgene plasmid # 52111)(Holst et al., 2006)
- OT-I TCR plasmids were deposited to <u>Addgene</u> by us:
  - pcDNA3.1(+)-OTI-TCRA_p2_TCRB (#131033)
  - pcDNA3.1(+)-OTI-TCRB_p2_TCRA (#131034)
  - pcDNA3.1(+)-OTI-TCRA (#131035)
  - pcDNA3.1(+)-OTI-TCRB (#131036)

### Plasmid labeling with Label-IT kit

100 ug of pCMV6 plasmid was labeled with 55 ul of Cy5 Label-IT kit for 1 hour at 37°C (Mirus Bio, Madison, WI). The labeled plasmid was purified by ethanol precipitation. In brief, 0.1 volume of 5M sodium chloride and 2.5 volumes of ice-cold 100% ethanol was added to the reaction. The solution was mixed and the tube was kept at −20°C for at least 30 minutes. Following the centrifugation and ethanol wash, the DNA pellet was resuspended in 10 mM Tris-Cl buffer (pH 8.5) and the DNA absorbance was read at A260 by NanoDrop One (Thermo Fisher) to quantify the eluted DNA.

### Staining and imaging of T cells

The cells were collected and centrifuged at 300 x g for 5 minutes. The supernatant was discarded and the cells were washed once with PBS. Then, the cells were resuspended in PBS and 16% formaldehyde (Thermo Fisher #28908) was added at a final concentration of 4%. The cells were fixed for 30 minutes at 4°C. After incubation, the cells were pelleted and washed twice with 1X BD Perm/Wash buffer (BD Biosciences #554714, Franklin Lakes, NJ). After the wash, the cells were stained with Alexa Fluor 488 Phalloidin (Thermo Fisher #A12379) for 30 minutes at room temperature in dark. After the incubation, the cells were pelleted and washed with PBS. In the end, the cells were resuspended in PBS and cytospinned on microscope slides (VWR, Radnor, PA, #48312-004) by centrifugation for 5 minutes at 500 x g. After the spin, 1 drop of ProLong Glass Antifade Mountant with NucBlue Stain (Thermo Fisher #P36983) was added on the slide and the cells were covered with a coverslip (size 22×22 mm). The cells were visualized by the Keyence BZ-X710 fluorescence microscope at 60X magnification (Nikon, Plan Apo, 60X/1.40 Oil; MRD01605) at room temperature. Filter cubes used: DAPI: OP-87762; GFP: OP-87763; TRITC: OP-87764

Protocol details: DOI:10.17504/protocols.io.vede3a6.

### Image analysis with Cytokit

Image analysis was conducted using Cytokit pipelines configured to segment nuclei over U-Net probability maps (McQuin et al., 2018) followed by secondary (cell boundary) and tertiary (plasmid body) object detection using threshold images resulting from Phalloidin and labeled plasmid channels. All image objects were subjected to morphological and minimum intensity filters before establish nucleus localization frequencies for plasmid objects, and parameters for this filtering were varied in a sensitivity analysis to ensure that findings are robust to processing configuration. Single-cell image visualizations were generated using Cytokit Explorer. Raw imaging data sets are publicly available at the following Google Storage URL: gs://cytokit/datasets/dna-stain.

### In vitro transcription (IVT)

IVT was performed using the T7 promoter of the plasmids and HiScribe T7 ARCA mRNA kit with tailing (NEB #E2060S, Ipswich, MA). The whole kit was used with 20 ug linearized DNA following the manufacturer’s protocol. The final RNA product was eluted in 330 ul nuclease-free water and incubated at 65C for 10 mins before measuring RNA concentration by the NanoDrop One.

### Electroporation of T cells

#### Human

After 2 days of activation, the cells were collected and put in a centrifuge tube. The tube was placed on DynaMag (Thermo Fisher) and the magnetic beads were removed. Activated and unstimulated cells were centrifuged for 7 minutes at 350 x g, the supernatant was aspirated and the cell pellet was washed once with PBS and then resuspended in electroporation buffer (R for activated cells, T for unstimulated cells) (Thermo Fisher). When working with Neon 10 ul tip, 200,000 cells were resuspended in 9 ul of T buffer and 1.5 ug DNA was added. Electroporation was performed at 1600 V 10 ms 3 pulses settings for activated cells and at both 2200 V 20 ms 1 pulse and at the same settings as activated cells for unstimulated cells using Neon electroporation device (Thermo Fisher).

For DNA electroporation experiments in activated cells, 5 reactions were seeded on a 24-well-plate (a total of 1 million cells) with 0.5 ml T cell media. For DNA electroporations in unstimulated T cells, Neon 100 ul tip was used and 2 million cells were electroporated per reaction and then plated on a 24-well-plate with 1 ml media.

For mRNA electroporations, cell pellet needs to be washed thoroughly with PBS to get rid of any potential RNases in the cell media. For mRNA electroporation of activated cells, Neon 100 ul tip was used and 1 million cells were electroporated per reaction and then plated on a 24-well-plate with 1 ml media and 200IU/ml IL-2. For OT-I TCR (mRNA) electroporation, 5 ug OT-I TCR alpha mRNA and 5 ug OT-I TCR beta mRNA were electroporated into 1 million activated cells. For mRNA electroporation of unstimulated cells, Neon 100 ul tip was used and 1-1.5 million cells were electroporated per reaction and then plated on a 24-well-plate with 1 ml media. For each mRNA electroporation reaction, one Neon 100 ul tip was used. For the microscope imaging experiment, 3x Neon 100 ul reactions (6 million cells and 45 ug labeled DNA in total) were electroporated and plated on a 12-well-plate with 3 ml T cell media.

#### Mouse

Activated WT Mouse CD8+ T cells were electroporated on the 4th or 5th day of activation based on their cell count. The cell pellet was collected by centrifugation at 450 x g for 5 mins. 2x PBS washes were performed before resuspending the cell pellet in R buffer (Neon systems, Thermo Fisher). 1-2 million cells were electroporated per reaction using the Neon electroporation device at 1300V 10ms 3 pulses setting. 10 ug (IVT’ed) GFP RNA was used for 1 million cells. For OT-I TCR mRNA electroporations, 5 ug OT-I TCR alpha mRNA and 5 ug OT-I TCR beta mRNA were used per million cells. For the single mRNA constructs (OT-I TCR alpha-linker-beta or beta-linker-alpha), 10 ug mRNA were electroporated into 1 million cells.

### Cas9 RNP preparations and electroporation

Cas9 RNPs were prepared immediately before the experiment. For activated cells, only one single crRNA (Thermo Fisher) was mixed with tracRNA (Thermo Fisher) and incubated at a thermocycler for 5 mins at 95C and 25 mins at 37C. After incubation, the newly formed sgRNA (7.5 pmol sgRNA for 200,000 cells) was mixed with TrueCut v2 Cas9 protein (0.25 ul of Cas9 for 200,000 cells; #A36499, Thermo Fisher) and incubated in the cell culture incubator (at 37C) for 15-20 mins. Then, the cells were added on top of the prepared Cas9 RNPs and immediately were electroporated. For the unstimulated cells, 3 crRNAs were used per target. Individual crRNAs were incubated with equal volumes of tracRNA. After thermocycler incubation of the individual sgRNAs was completed, 3 sgRNAs were mixed together and then Cas9 protein was added. The protocol for Cas9 RNP preparation for unstimulated cells and the crRNA sequences for CXCR4 and CD127 were adapted from Seki and Rutz (Seki and Rutz, 2018).

### CD8 depletion of unstimulated T cells

Dynabeads™ Pan Mouse IgG beads (Thermo Fisher #11041) were used with purified CD8 antibody (Biolegend, San Diego, CA) to deplete CD8+ cells from unstimulated pan T cells. The manufacturer’s protocol for the “indirect technique” was followed. Depletion efficiency was checked by the flow cytometer.

### CD8 enrichment of human and mouse T cells

The CD8+ T cell enrichment was performed either right after debeading the activated cells on day 2 for human cells and day 3 for mouse cells or the next day following the debeading. Human CD8+ T cells were enriched by Dynabeads™ Untouched™ Human CD8 T Cells Kit (Thermo Fisher #11348D) and mouse CD8+ T cells were enriched by Dynabeads™ Untouched™ Mouse CD8 Cells Kit (Thermo Fisher #11417D). The manufacturer’s protocols were followed.

### crRNA sequences

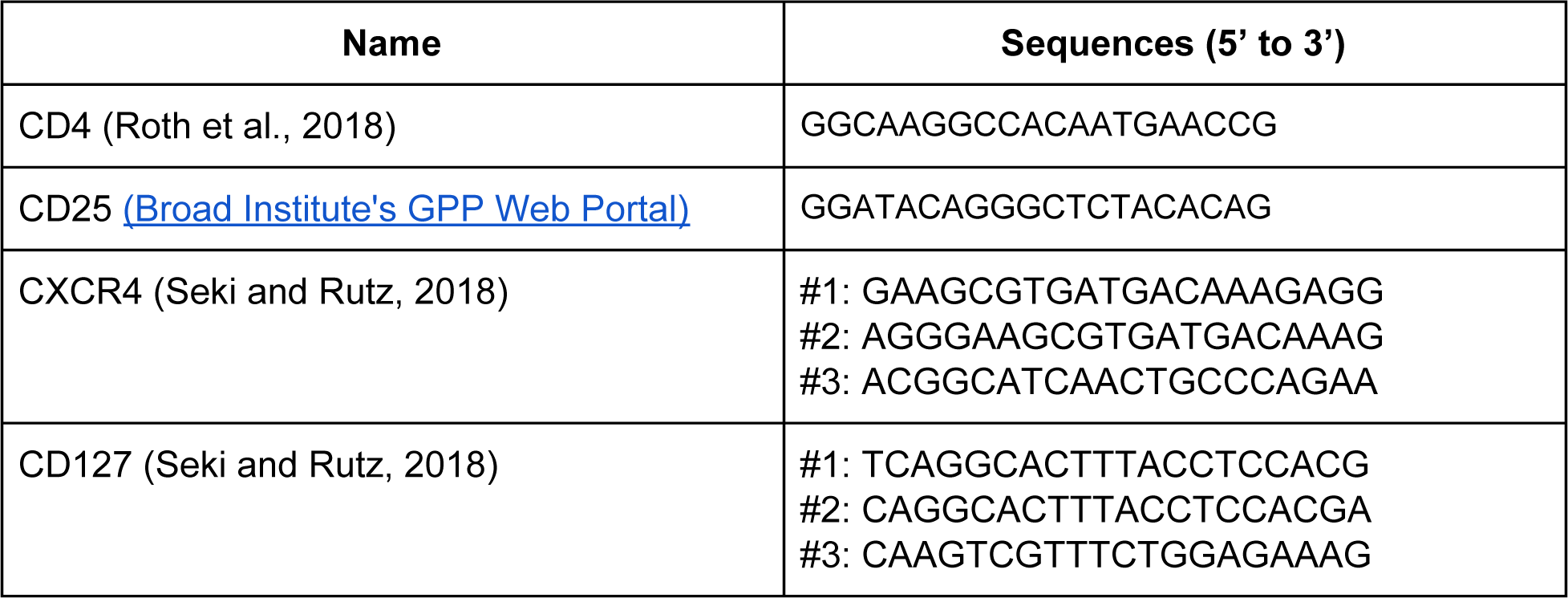

### Antibodies

| Name | Vendor | Catalog # |
| --- | --- | --- |
| CD45RO | Biolegend | 304210 |
| CCR7 | Biolegend | 353212 |
| CXCR4 | Biolegend | 306518 |
| CD127 | Biolegend | 351310 |
| CD4 | Biolegend | 317418 |
| CD25 | Biolegend | 302627 |
| CD8 | Biolegend | 344710 |
| CD8 (mouse) | Biolegend | 100734 |
| CD8 (depletion) | Biolegend | 344702 |

### H2Kb-SIINFEKL Multimers

| Name | Vendor | Catalog |
| --- | --- | --- |
| H-2Kb SIINFEKL | Immudex | JD2163 (APC) |
| H-2Kb SIINFEKL | Tetramer Shop | MKb-001 (APC) |

### Flow Cytometry

Flow cytometry was performed on BD FACSVerse Flow Cytometer. Cells were collected and centrifuged at 300 x g for 5 minutes. The cell pellet was resuspended in flow buffer (PBS with %20 FBS) and the labeled-antibodies were added at the recommended concentration. The cells were stained at room temperature for 20-30 minutes in the dark. After incubation, the cells were pelleted and resuspended in PBS.

For the multimer-staining of T cells, 300,000 cells were collected in each microcentrifuge tube. The cells were centrifuged at 300 x g for 5 minutes. The cell pellet was resuspended in 50 ul flow buffer and 5 ul of the H2Kb-SIINFEKL-dextramer (Immudex, Copenhagen, Denmark) or 5 ul of the H2Kb-SIINFEKL-tetramer (Tetramer Shop, Kongens Lyngby, Denmark). The cells were incubated with the multimers for 10 mins at room temperature in the dark. Following that, 50 ul flow buffer and 2 ul CD8 antibodies were added and the cells were incubated for another 20 mins at room temperature in the dark. After incubation, the cells were washed with PBS, pelleted, and finally resuspended in PBS. Flow cytometry results were analyzed by FlowJo v10 (TreeStar, Ashland, OR, USA). The graphs were generated using GraphPad Prism8 software (GraphPad Software, San Diego, CA, USA).

### Cytotoxicity Assays

#### Cytotoxicity assay by LDH-release method

To estimate the T cell-mediated killing of the MC38 cells, we used CytoTox-ONE Homogeneous Membrane Integrity Assay (Promega, Madison, WI). Using a 96-well-plate, starting from the second column and second row, 25,000 MC38s were plated in 100 ul complete DMEM to fill 36 wells (6 wells in each row and column). For each different T cell population, 2x 96-well-plates were prepared (one plate for pulsing with the peptide and the other plate to keep the MC38 cells as unpulsed). The next day, T cells were centrifuged and their concentration was adjusted to 2 million cells per ml with 200IU/ml IL-2. For the plate that was going to be pulsed, the SIINFEKL peptide (Genscript #RP10611, Piscataway, NJ) was added to the T cell stock (10 uM) and also to the T cell media that was going to be used for the serial dilutions. Media was aspirated from the wells and 100 ul T cells were directly added to the 2nd and the 8th column. 100 ul T cell media was added into the columns 3-7 and 9-12. 100 ul T cells were again seeded from the stock solution to the 3rd and 9th column and from there they were serially diluted into the 4th and 10th and eventually to the 5th and 11th column. The extra 100 ul was discarded from columns 5 and 11. This way, column 2 had 4:1 E:T ratio, column 3 had 2:1, column 4 had 1:1, and column 5 had 1:2. Columns 6 and 7 had only MC38s. Columns 8-11 had only T cells and column 12 had only T cell media to control for the background signal. Cells under column 7 were lysed using the lysis buffer provided in the kit and the signal from this column was assumed to be the maximum LDH signal. We followed the manufacturer’s protocol for performing the assay and the plates were read by SpectraMax i3x (Molecular Devices, San Jose, CA) at ex/em 560nm/590nm. This protocol was adapted from http://dx.doi.org/10.17504/protocols.io.6j4hcqw.

#### Cytotoxicity assay by flow cytometry

Protocol details: http://dx.doi.org/10.17504/protocols.io.8d2hs8e

## Data Availability

Intermediate and final data sets that were used to generate the figures and summaries in this manuscript are available at https://github.com/hammerlab/t-cell-electroporation.

## Author Contributions

PA led the study design and performed the experiments with assistance from BAA. EC analyzed the fluorescent microscopy images. JH supervised the study. PA wrote the manuscript and all other authors provided feedback on the manuscript.

## Acknowledgments

The authors would like to thank Alexander Marson and Theo Roth for their initial guidance on the use of Cas9 RNPs in primary T cells; Paulos Lab and Rubinstein Lab for their feedback on the manuscript and the project; Mehrotra Lab for their help with cytospin. This work is supported in part by the Flow Cytometry and Cell Sorting Unit Shared Resource, Hollings Cancer Center, Medical University of South Carolina (P30 CA138313). The authors declare no competing financial interests.

## Supplemental Data

**Figure S1.**
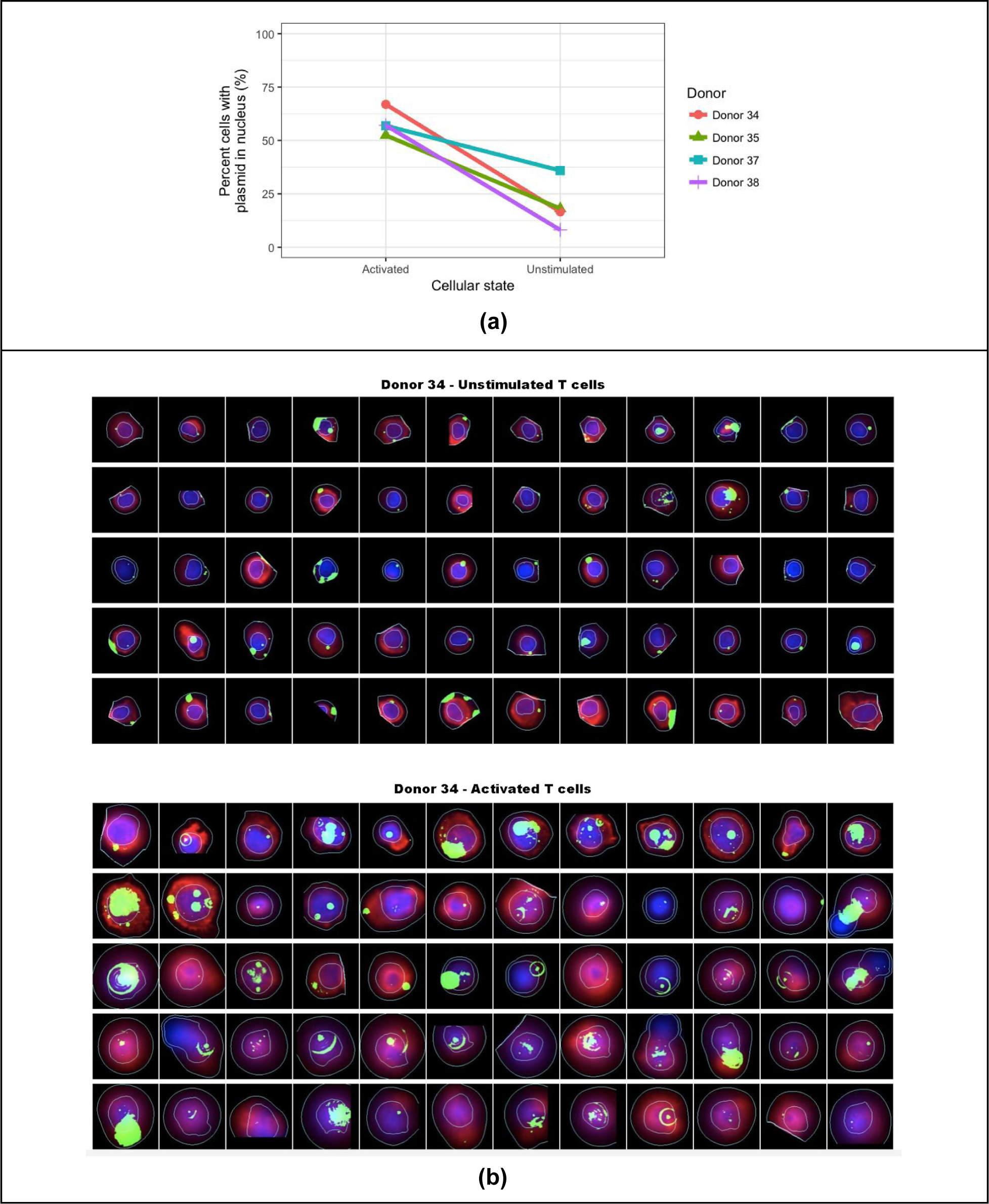
Imaging of the labeled plasmid electroporated cells 24 hours after electroporation. Cy5-labeled-pCMV6 was electroporated into unstimulated and activated cells on the second day of activation. The cells were incubated for 24h and then they were fixed on slides. (a) The frequency of cells with nuclear plasmid was higher in activated T cells compared to unstimulated T cells. See <u>this notebook</u> for detailed data and analysis. (b) Cytokit was used to analyze the microscope images (Czech et al., 2018). Each sub-panel shows 60 representative individual cells that were plasmid positive. Cell, nucleus, and plasmid signal borders as well as the signal intensities are shown as inferred via Cytokit’s detection algorithm (Red: phalloidin, green: Cy5-labeled-plasmid, blue: DAPI). See Table S1 for detailed inferred cellular characteristics.

**Table S1.**
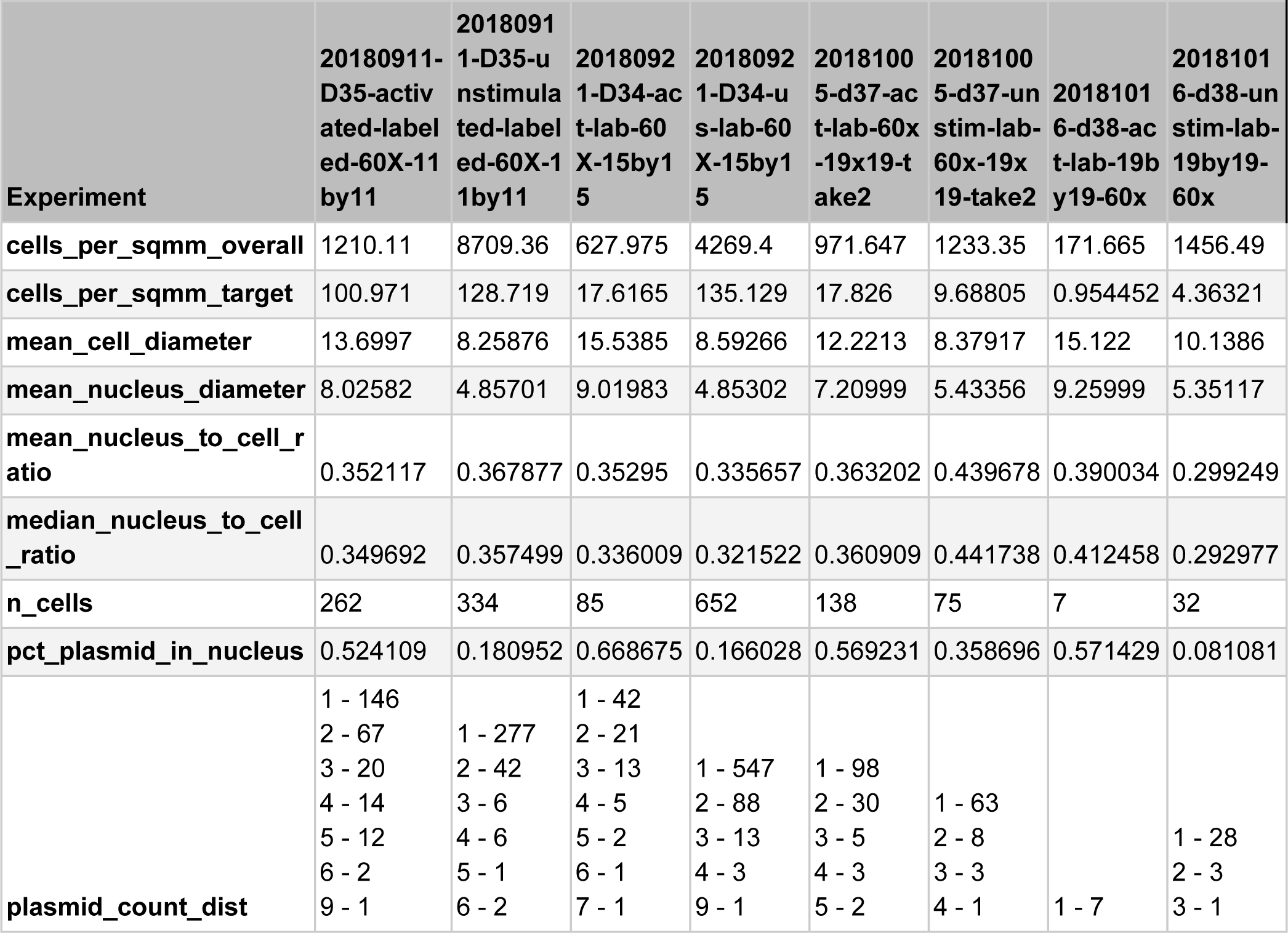
Details of cellular characteristics inferred from fluorescence microscopy images of activated and unstimulated cells via Cytokit.

**Figure S2.**
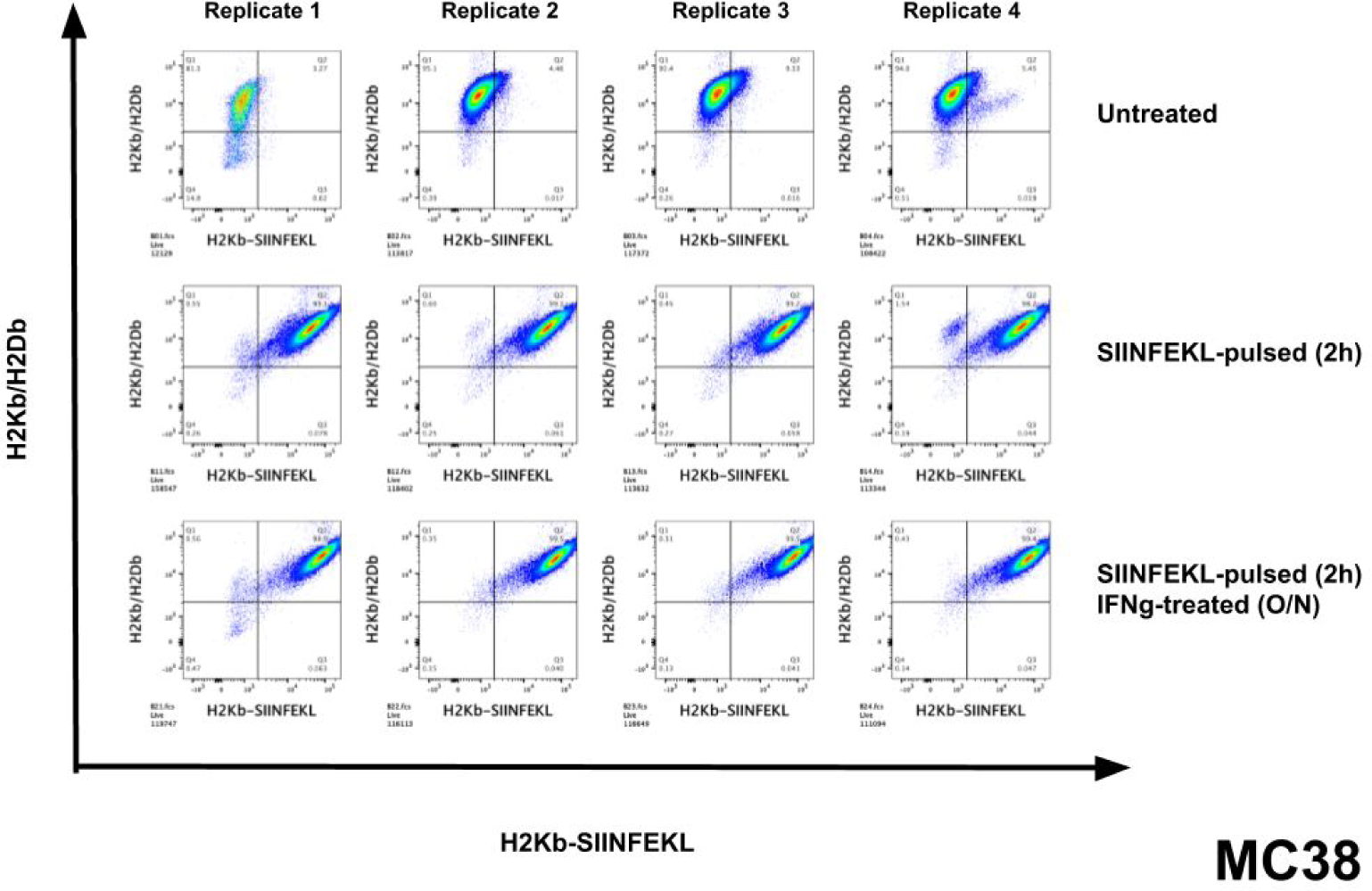
SIINFEKL presentation by MC38 cells. When the cells were untreated there was no staining by the H2Kb-SIINFEKL multimer as expected (a), however when the cells were pulsed with the peptide for 2hrs, almost all of the cells presented the peptide (b) and the presentation was further enhanced with a prior overnight IFN-g (10 ng/mL) treatment.

**Figure S3.**
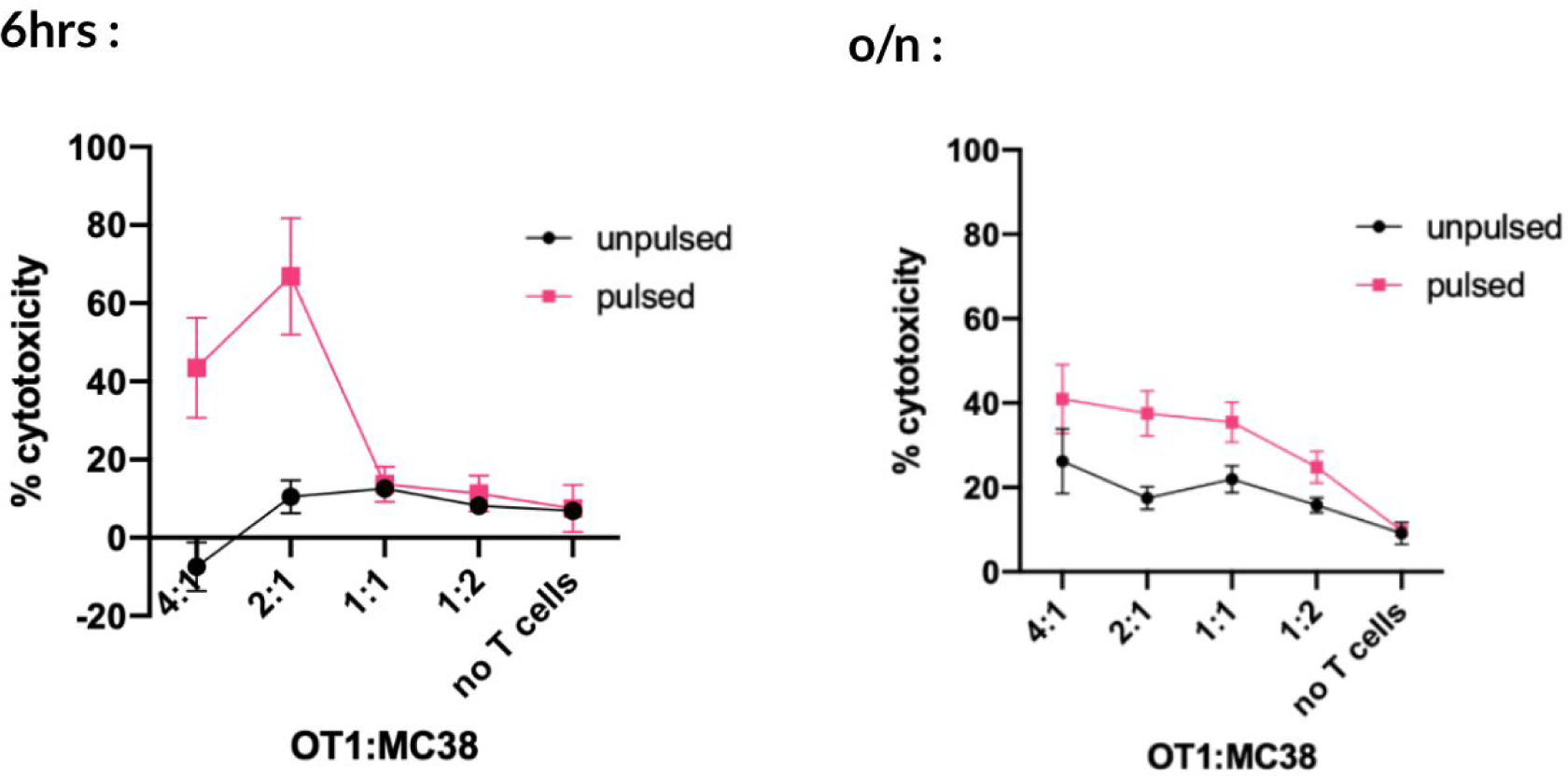
Cytotoxicity assay (LDH-release) for OT-I cells and SIINFEKL pulsed or unpulsed MC38s. The co-cultures with varying OT-I to MC38 ratios were either incubated for 6hrs or overnight and the cytotoxicity assay was performed. 6-hour co-culturing was able to capture the cytotoxicity activity better compared to the overnight co-culturing.

**Figure S4.**
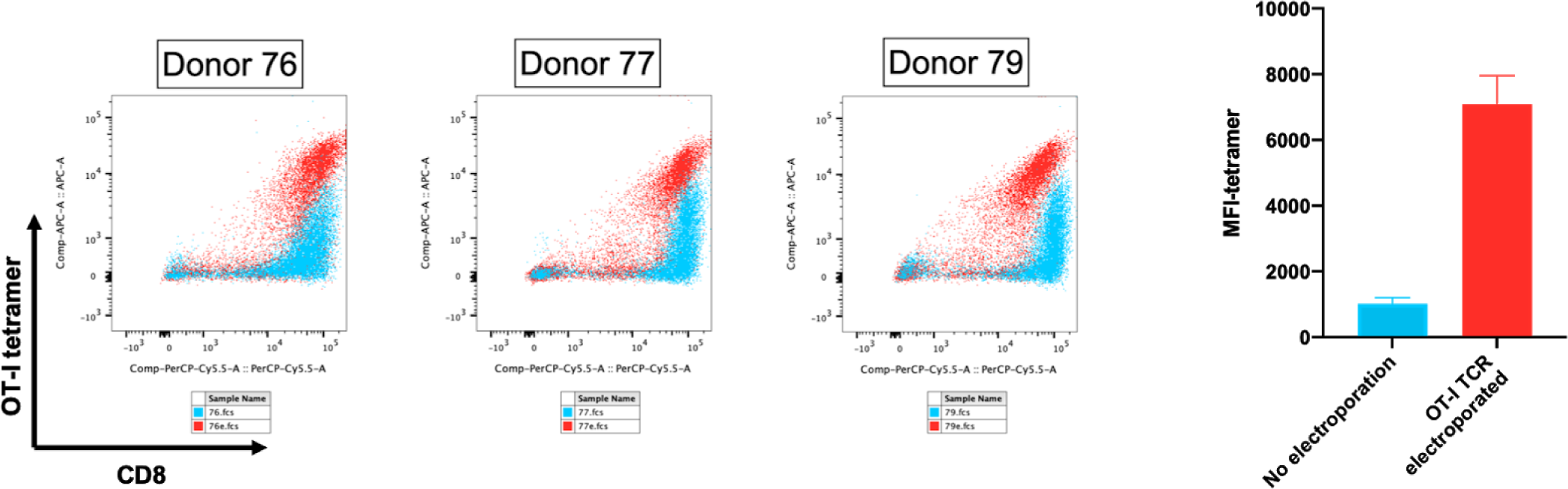
Multimer staining of OT-I TCR electroporated human CD8+ T cells. CD8+ T cells from 3 donors were electroporated with OT-I TCR alpha and beta mRNAs. The next day, electroporated (red) and unelectroporated (blue) cells were collected and stained with H2Kb-SIINFEKL multimer and CD8 antibody. The samples were analyzed by flow cytometry and the flow plots were generated by FlowJo v10.

